# Design of solubly expressed miniaturized SMART MHCs

**DOI:** 10.1101/2025.03.14.643101

**Authors:** William L. White, Hua Bai, Chan Jhong Kim, Kevin M. Jude, Renhua Sun, Laura Guerrero, Xiao Han, Xiaojing Tina Chen, Apala Chaudhuri, Julia E. Bonzanini, Yi Sun, Amarachi E. Onwuka, Nan Wang, Chunyu Wang, Xinting Li, Inna Goreshnik, Aza Allen, Paul M. Levine, Hao Yuan Kueh, Michael C. Jewett, Nikolaos G. Sgourakis, Adnane Achour, K. Christopher Garcia, David Baker

## Abstract

The precise recognition of specific peptide-MHC (pMHC) complexes by T-cell receptors (TCRs) plays a key role in infectious disease, cancer and autoimmunity. A critical step in many immunobiological studies is the identification of T-cells expressing TCRs specific to a given pMHC antigen. However, the intrinsic instability of empty class-I MHCs limits their soluble expression in *Escherichia coli* (*E. coli*) and makes it very difficult to characterize even a small fraction of possible pMHC/TCR interactions. To overcome this limitation, we designed small proteins which buttress the peptide binding groove of class I MHCs, replacing β2-microglobulin (β2m) and the heavy chain α3 domain, and enable soluble expression of both H-2D^b^ and A*02:01 in *E. coli*. We demonstrate that these soluble, monomeric, antigen-receptive, truncated (SMART) MHCs retain both peptide- and TCR-binding specificity, and that peptide-bound structures of both allomorphs are similar to their full-length, native counterparts. With extension to the majority of HLA alleles, SMART MHCs should be broadly useful for probing the T-cell repertoire in approaches ranging from yeast display to T-cell staining.

**Significance:** Despite the critical role that TCR/pMHC interactions play in human health, it has remained difficult to produce reagents necessary to study them. Requirements for refolding or sequence optimization limit immunologists’ and biochemists’ ability to characterize diverse pMHC/TCR interactions. Here, we develop a *de-novo* designed protein domain that stabilizes the H-2D^b^ and A*02:01 class I MHC allomorphs, allowing soluble expression in *E. coli* without the need for a stabilizing peptide, and improving display on the yeast surface, while maintaining peptide and TCR binding interactions. These features facilitate a wide range of experiments to more fully understand the nature of pMHC/TCR interactions, and pave the way for the development of stabilizing domains for all MHC allomorphs.

## Introduction

Recombinantly expressed peptide-major histocompatibility complexes (pMHCs) are widely used as staining reagents to identify or isolate T-cell or NK-cell subsets that recognize a peptide of interest^1^. They have been widely used to study T-cell specificity^2–4^, infectious disease^5^, autoimmunity^6,7^, and cancer immunology^8–10^. Recombinant pMHCs have also been critical in determining the structures of peptide/MHC and pMHC/T-cell receptor (TCR) complexes^2,11,12^. All of these discoveries were made despite the significant difficulties involved in producing the soluble pMHCs that are necessary for the underlying biophysical, structural, and functional experiments. The pMHC production process, which involves separate expression in *E. coli* of each of the two MHC chains as insoluble inclusion bodies, solubilization, and subsequent refolding in the presence of the desired peptide^13^, is expensive, slow, and inefficient. To reduce the burden of refolding, systems have been developed in which a single refolding reaction can be split and loaded with many different peptides^14–17^; these methods have increased the number of peptide variants that can be studied, but are still limited by the need for refolding. Eukaryotic expression systems that fold the MHC structure natively^18–22^ have been used in peptide library screens where peptide variants are fused to the MHC and displayed on the surface of a cell, but can be limited in library size or expression levels.

The difficulties in producing pMHCs likely stem from the inherent instability of the MHC molecule when either a peptide or the β2m subunit is absent. A more stable MHC-like molecule that could be readily expressed in *E. coli* or yeast would enable the study of peptide-specific T-cell populations at a much larger scale. Instead of relying on specialized facilities to produce refolded pMHCs for staining experiments^13^, immunologists could produce them in-house, dramatically improving their ability to iterate through multiple peptide variants or MHC alleles. Similarly, a stabilized native-like MHC could facilitate screening of large peptide libraries by yeast display without the need for prior optimization of the MHC sequence for display. We reasoned that such a molecule could be created by leveraging the stability of *de-novo* designed proteins and recent advances in protein-protein interface design^23^ to replace portions of the MHC molecule with a designed protein scaffold. We set out to design soluble, monomeric, antigen-receptive, truncated (SMART) MHC molecules that replace both the α3 domain and the β2m subunit with a small designed protein domain that buttresses the peptide binding groove of the pMHC. These stabilizing domains should preserve the native peptide- and TCR-binding properties, and allow soluble expression in the absence of a peptide without the need for refolding, providing a peptide-receptive MHC that can be loaded with arbitrary peptides.

### Design of SMART MHCs

The structure of a native class I MHC is composed of a heavy chain, the β2m subunit, and a peptide^24^. The α1 and α2 domains of the heavy chain consist of a β-sheet supporting two α-helices which create a peptide-binding groove that defines the peptide binding specificity of each MHC-I allomorph and facilitates interactions with TCRs^24^. The α3 domain of the heavy chain, and the β2m subunit, are more membrane proximal and function as structural support for the α1 and α2 domains^24^. The α3 domain additionally provides a binding site for the CD8 co-receptor on T-cells^25^. Since all class I pMHCs share very similar structures, we selected the mouse H-2D^b^ allele, to use as a representative in our stabilizing design process (fig. 1A).

**Figure 1.**
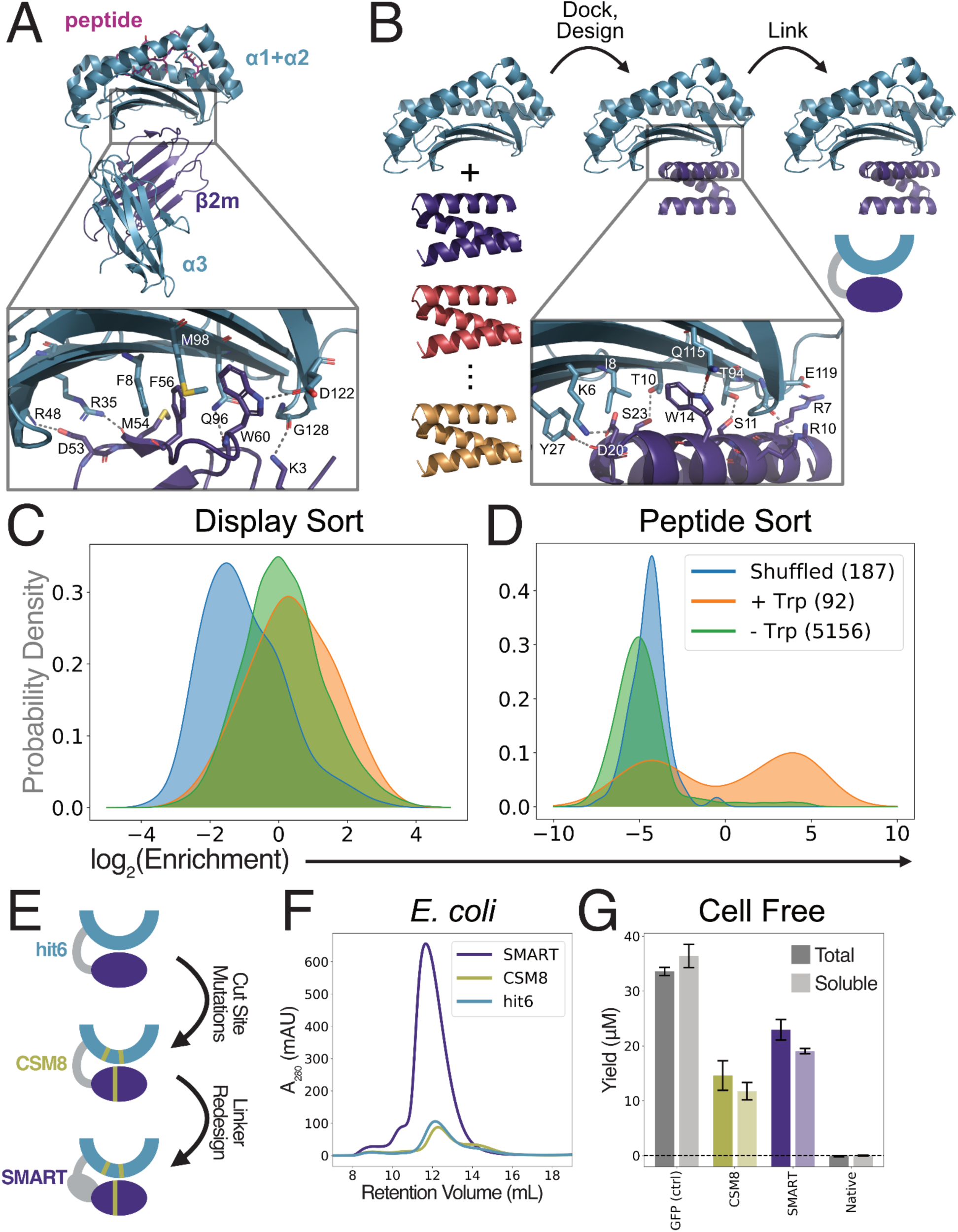
Design and screening of SMART MHCs. A) Structure of native H-2D^b^/β2m/gp33 complex. Inset shows selected interactions that are formed between the β2m subunit and the α1 and α2 domains. B) Schematic of the initial design process used to generate a library of candidate stabilizing domains showing the truncated H-2D^b^ with candidate stabilizers (left), the placement of the stabilizing domain (middle), and the final linked version used for yeast display (right). Inset shows selected interactions between the stabilizing domain and the α1 and α2 domains. C-D) Kernel density estimates (KDEs) of the distribution of enrichment values of all designs in either the expression (C) or peptide binding (D) yeast display sorts. Designs are split into three categories for KDE calculations: sequence-shuffled negative controls (shuffled), designs that do not replace W60 from β2m (-Trp), and designs that do replace W60 from β2m (+Trp). Relative enrichment for each design was calculated by dividing its abundance after the indicated sort by its abundance prior to that sort. E) Schematic of the iterative improvements from hit6 to the final SMART design, showing the cleavage site mutations and the redesigned linker. F) Size exclusion chromatograms of soluble material from *E. coli* expression cultures. G) Total and soluble yields from cell free expression reactions for GFP (positive control) and different H-2D^b^ variants.

We began by removing as much of the structure as possible without disrupting the ability of the MHC to present peptides and interact with TCRs. Following the precedent of previously published truncated “mini-MHC” systems, we removed the α3 and β2m domains completely^11,26,27^ (fig. 1B, left). The truncated MHC heavy chain (residues 1-179) contains a relatively hydrophobic patch on the underside of its β-sheet which we used as the starting point for our stabilizing domain design. We used methods developed for the design of protein binders^28^ to design this *de-novo* stabilizing domain. First, we collected a set of candidate backbone structures for the stabilizing domain (fig. 1B, left). Next, we docked these backbones against the truncated H-2D^b^ structure and designed their sequences to create favorable contacts with the MHC, replacing the interactions made by the deleted α3 domain and β2m subunit (fig. 1B, middle). Finally, we linked each designed stabilizing domain to the N-terminus of the truncated MHC by a poly-GGS linker (fig. 1B, right).

We screened a total of about 10^4^ designs generated by this method using yeast surface display^29^. We first sorted for designs that enabled the display of truncated H-2D^b^ on the yeast surface (fig. S1A), and within that sorted population, we further sorted for designs which were able to bind to a FITC-labeled gp33 peptide (FITC-gp33) (fig. S1B), which is bound strongly by native H-2D^b^ ^30,31^. The display sort provided only slight separation of our designs from sequence-scrambled negative controls, while the binding sort allowed us to clearly distinguish successful designs from controls (fig. 1C-D). We found that designs which contained a Trp residue placed similarly to W60 of β2m in the native H-2D^b^/β2m/gp33 structure were positively enriched, which aligns well with previous observations that mutations at W60 significantly destabilize the interaction between β2m and the MHC heavy chain^32,33^.

We identified the 30 designs with the highest peptide-binding enrichment and expressed them in *E. coli.* The best performing of these stabilizing domains (hit6; fig. 1E, top) enabled partial MHC folding in the absence of a peptide in both yeast and *E. coli* expression systems. However, in *E. coli* only a fraction of the hit6 protein was soluble, and was susceptible to proteolysis, resulting in very low yields (fig. 1F, teal curve). To improve hit6, we identified likely cleavage sites based on the masses of the proteolytic fragments, and redesigned the amino acid sequence near those sites, keeping the amino acids that directly interact with the peptide or TCR fixed, and biasing towards amino acids that occur frequently in other MHC alleles (fig. S2). We chose the cleavage site mutant (CSM) with the lowest degree of proteolysis, called CSM8 (fig. 1E, middle), and sought to further improve it by incorporating a more rigid linker to reduce domain-swapping and off-target misfolding. We used inpainting, ProteinMPNN, and AlphaFold2^34–36^ to design such a linker, replacing the flexible GS linker in hit6 and CSM8 (fig. 1E, bottom). We refer to the most soluble of these improved designs as SMART MHC throughout the remainder of the text. The results presented below make use of all three design variants because some experiments were performed prior to the development of the SMART variant, and because the CSM8 mutations are not necessary in some experimental contexts. To distinguish between these variants, data in all figures are colored to match each design variant (hit6: teal; CSM8: yellow; SMART: purple). We tested the expression of the SMART constructs in a cell-free expression (CFE) system which is compatible with a variety of high-throughput screening methods^37–42^. Engineered CFE systems can produce high protein yields^43^ and create oxidizing environments for disulfide bond formation, critical for maintaining structural integrity of eukaryotic proteins such as MHCs^44–46^. The cell-free results confirmed the *E. coli* results, demonstrating that, in contrast to the native H-2D^b^, the stabilized designs can be expressed solubly, and that SMART H-2D^b^ is expressed more efficiently than CSM8 H-2D^b^ (fig. 1G). These results indicate that SMART MHCs can be expressed in a variety of systems that are not compatible with soluble expression of native MHC-I molecules.

### SMART H-2D^b^ retains native binding properties and structure

To verify that the soluble, peptide-free SMART H-2D^b^ material produced in *E. coli* retained the functional characteristics of the native MHC, we first measured the binding affinity of SMART H-2D^b^ for FITC-gp33 using fluorescence polarization (FP). These measurements demonstrated that SMART H-2D^b^ binds FITC-gp33 with an apparent dissociation constant (K_D,app_) below 1nM (fig. 2A); this value islower than the previously reported value of 21nM for native H-2D^b^ ^47,48^, likely because competitor peptide was included in the native measurement. SMART H-2D^b^ was highly shelf-stable, retaining this strong peptide binding affinity even after storage for at least one month at 4°C, with only a minor decrease in affinity following multiple freeze/thaw cycles (fig. S3A-B). To further assess peptide binding and stability, we measured circular dichroism (CD) spectra and melting curves for SMART H-2D^b^ in the presence and absence of the gp33 peptide. The CD spectra are consistent with the mixed αβ fold of the design model (fig. S3C), while the melting curves demonstrate a clear stabilization of the fold in the presence of the peptide (fig. S3D).

**Figure 2.**
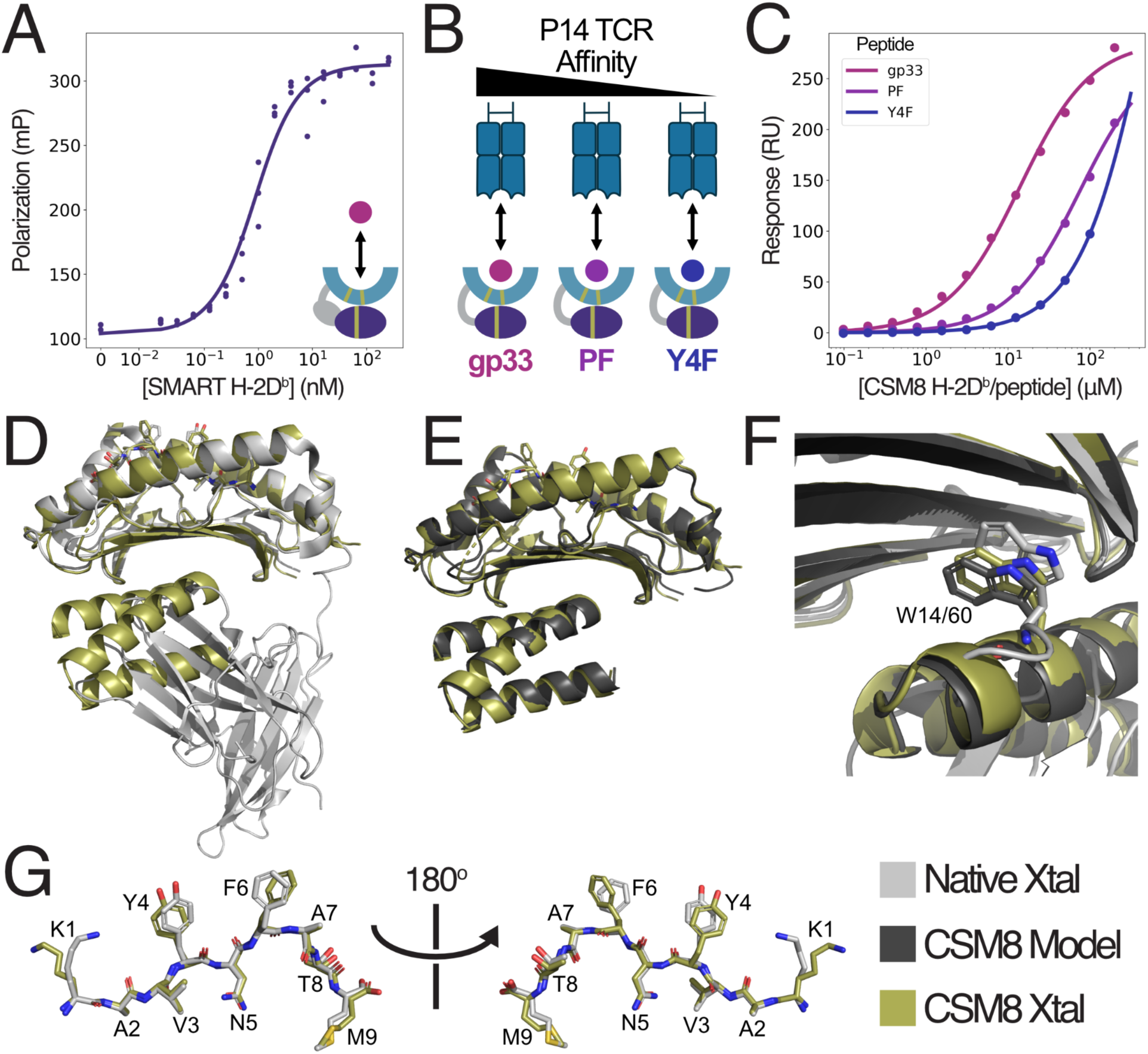
SMART H-2D^b^ closely recapitulates the structure and interactions of the native MHC peptide binding domain. A) FP data (filled circles) and fitted binding curves (lines) for FITC-gp33 binding to purified SMART H-2D^b^. Each point represents one of n=3 technical replicates. B) Cartoon representations of gp33 peptide variants with varying affinity for the P14 TCR. C) SPR equilibrium binding measurements (filled circles) and fitted binding curves (lines) for complexes of CSM8 H-2D^b^ with three different peptides binding to immobilized P14 TCR. D) Structural superposition of the crystal structure of the CSM8 H-2D^b^/gp33 complex (yellow; PDB ID: 9HY4) onto the previously determined H-2D^b^/β2m/gp33 complex (light gray; PDB ID: 1S7U). E) Structural alignment of the CSM8 H-2D^b^/gp33 complex crystal structure (yellow) to the design model (dark gray). F) Structural alignment of all structures from D and E highlighting the similar positioning of W60 in the native structure relative to W14 of the CSM8 stabilizing domain. G) Comparison of the conformation of the gp33 peptide in the two crystal structures.

Next, we assessed the binding affinity of the H-2D^b^/gp33-specific P14 TCR to CSM8 H-2D^b^ loaded with three variants of the gp33 peptide that bind to H-2D^b^ with similar affinity but altered recognition by the P14 TCR^30,49^ (fig. 2B). Surface plasmon resonance (SPR) measurements revealed TCR binding affinities similar to native H-2D^b^/peptide complexes for all three variants^49^ (table 1; fig. S4; fig. 2C), further indicating that our stabilizing domain maintains the peptide-binding domain and peptide in a native-like conformation.

**Table 1.**
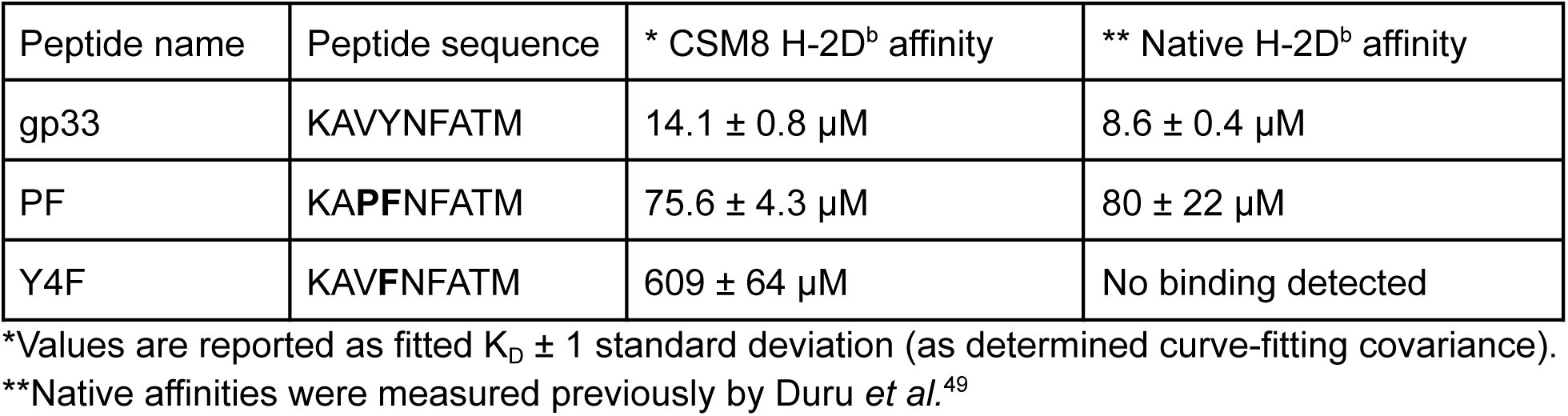
P14 TCR binding affinities of CSM8 H-2D^b^ in complex with three gp33 peptide variants. CSM8 affinities are calculated by fitting a standard binding curve to the equilibrium binding SPR values. Mutations relative to the gp33 sequence are shown in bold.

As a final validation of our design, the crystal structure of the CSM8 H-2D^b^/gp33 complex was determined to 2.0Å resolution. This high resolution allowed us to both analyze the interactions formed between the peptide binding cleft and the stabilizing domain underneath, as well as to compare the conformation of the presented peptide, and of residues known to be essential for TCR recognition, with the native H-2Db molecule presenting the same epitope^31,50^. The peptide binding groove of the CSM8 H-2D^b^ structure aligns very well with the native structure, with a Cα-RMSDs of 0.60Å over 176 atoms (fig. 2D). The structure of the stabilizing domain closely matches our computational design model, with a Cα-RMSD of 0.70Å over 233 atoms (fig. 2E). Examination of the region surrounding the β2m tryptophan residue W60 revealed that the Trp sidechain takes a similar conformation in the CSM8 design model and both crystal structures (fig. 2F). Furthermore, the peptide backbone is very well aligned with the native conformation, and the side chains display minor changes between the two crystal structures (fig. 2G; residues p1K, p4Y and p6F). These small changes, along with a few minor shifts elsewhere in the CSM8 H-2D^b^ structure could explain the slight deviations in peptide binding affinities we observed in our SPR experiments. Overall, these results demonstrate that SMART H-2D^b^ retains all the structural features that allow peptide and TCR binding, without the refolding requirements of the native MHC.

### The stabilizing domain allows soluble expression of functional HLA A*02:01

Next, we tested the ability of our stabilizing domain to generalize to other MHC allomorphs. Based on the structural and sequence similarity of H-2D^b^ to a variety of human MHCs we did not introduce any modifications to the stabilizing domain and instead focused on varying the MHC sequence. We selected HLA A*02:01 due to its high frequency across different ethnic groups^51^, and the large number of well-characterized peptide epitopes and TCRs that bind to this HLA allpmorph^52,53^. Although the CSM8 A*02:01 construct was easily expressed in a soluble form in *E. coli*, roughly half of the soluble material was in a dimeric (and likely misfolded) state, even after isolation of the monomeric fraction (fig. 3A). It should be noted that the monomer fraction was purified for use in the experiments described below.

**Figure 3.**
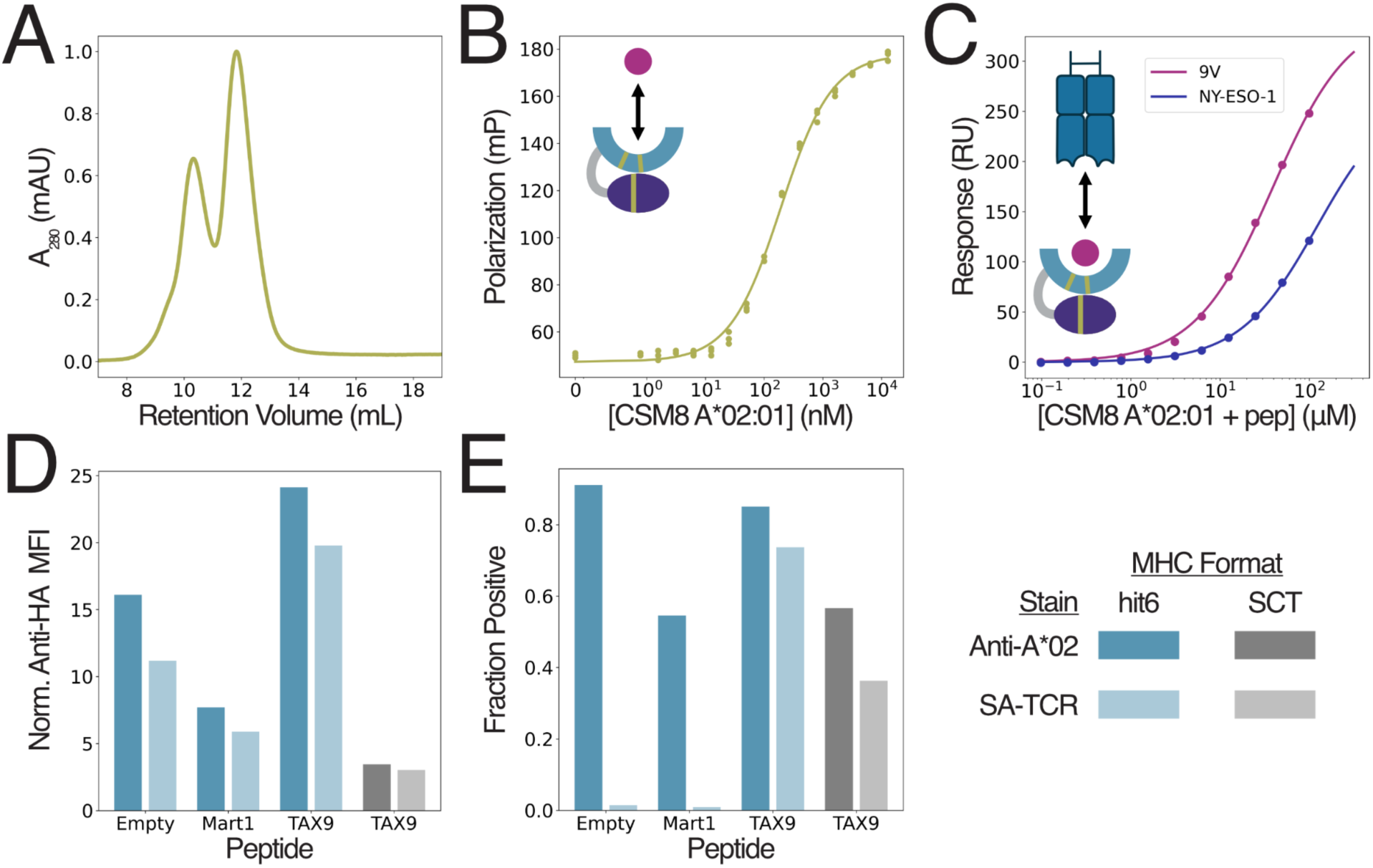
CSM8 A*02:01 retains native TCR interactions and improves pMHC yeast display. A) Size exclusion chromatogram trace of empty CSM8 A*02:01 purified from the soluble fraction of *E. coli* and thereafter from the monomeric fraction by SEC. B) FP data (filled circles) and fitted binding curve (line) for n=3 technical replicates of CSM8 A*02:01 binding to the AF488-NY-ESO-1 peptide. C) Equilibrium binding data (filled circles) and fitted binding curves (lines) from SPR experiments for CSM8 A*02:01 presenting variants of the NY-ESO-1 peptide to immobilized 1G4 TCR. D-E) yeast display data for hit6 (blue) or full-length SCT (gray) A*02:01 stained with an anti-HA antibody and either an anti-A*02 antibody (dark) or streptavidin A6c134 TCR tetramers (light). D) Surface expression levels (measured as anti-HA MFI) for the HA+ population of each sample. E) Fraction of HA+ yeast cells that are also stain+ for the anti-A*02 or TCR tetramer stain in each sample.

To test the ability of CSM8 A*02:01 to present peptides, we selected the tumor-associated antigen, NY-ESO-1^54^ and the well-characterized NY-ESO-1/A*02:01-specific TCR, 1G4^53^. We used FP methods very similar to those used to evaluate SMART H-2D^b^ to measure the K_D,app_ of NY-ESO-1 to CSM8 A*02:01 and found it to be approximately 190nM (fig. 3B). This value is somewhat weaker than the 40nM affinity measured for the native A*02:01 using competition binding assays^55^ which likely overestimate the K_D,app_, suggesting that the true difference may be larger. The difference in affinity between the native and CSM8 A*02:01 may reflect the presence of misfolded monomers, or re-equilibration to a monomer/dimer mixture, lowering the effective concentration of the monomer.

Given that CSM8 A*02:01 can bind the NY-ESO-1 peptide, albeit with reduced affinity, we next tested whether it also retained binding to the 1G4 TCR^54^. Using similar SPR methods to those used for H-2D^b^ we determined the affinity of the NY-ESO-1/CSM8 A*02:01 complex to the 1G4 TCR with two variants of the peptide (fig. 3C). We found that the measured affinities were roughly 7-9 fold weaker than the same values for native A*02:01 (table 2). Despite this difference in absolute binding affinity, the ranking of the affinities for peptide variants was maintained (table 2; fig. 3C). Overall, these data suggest that our designed domain can improve the stability of A*02:01, without the need for extensive redesign or screening.

**Table 2.** 1G4 TCR binding affinities for CSM8 A*02:01 in complex with NY-ESO-1 peptide variants. CSM8 affinities are calculated by fitting a standard binding curve to the equilibrium binding SPR values. Mutations relative to the NY-ESO-1 sequence are shown in bold.

| Peptide name | Peptide sequence | * CSM8 A*02:01 affinity | ** Native A*02:01 affinity |
| --- | --- | --- | --- |
| NY-ESO-1 | SLLMWITQC | 123 ± 6.1 µM | 13.3 ± 0.4 µM |
| NY-ESO-1-9V | SLLMWITQ**V** | 38.9 ± 2.0 µM | 5.7 ± 0.2 µM |

\*Values are reported as fitted $K_D$ ± 1 standard deviation (as determined curve-fitting covariance).
\*\*Native affinities were measured previously by Chen *et al*<sup>63</sup>.

As noted above, yeast display can be a powerful tool to screen libraries of peptide variants for TCR binding^19,22^, and we next evaluated the ability of SMART A*02:01 to express on the yeast surface, a necessary pre-requisite for library screening. We used hit6 A*02:01 for these experiments as the CSMs are not necessary in yeast display, and fused the TAX9 peptide^56^ to it, including the W167A MHC mutation to allow for the presence of a peptide linker^19^ (fig. S5B). We compared hit6 A*02:01 to native A*02:01 single-chain trimers (SCT) (fig. S5A) and found that it is displayed at much higher levels, regardless of whether a peptide is fused to it (fig. 3D). The elevated expression levels also led to an increase in binding of TCR tetramers made using the high-affinity A6c134 TCR^52,56^ when the TAX9 peptide was present, but not with an empty MHC or an unrelated control peptide (fig. 3E; fig. S5C-D). Thus, hit6 A*02:01 significantly enhances yeast display of HLA A*02:01 while retaining TCR binding specificity. These results indicate that hit6 A*02:01 could facilitate large peptide-MHC library screens.

### CSM8 A*02:01 presents TAX9 peptide to a TCR in a native-like manner

To investigate whether SMART A*02:01 presents peptides and interacts with TCR in a native-like way, we determined the crystal structure of CSM8 A*02:01/TAX9 complexed with the A6c134 TCR. Refolding methods were necessary to make sufficient quantities of monomeric CSM8 A*02:01 for crystallography. We found excellent agreement between CSM8 and native A*02:01 within the HLA (Cα-RMSD 0.40 Å over 161 atoms) and TAX9 peptide (all-atom RMSD 0.51 Å) (fig. 4A). CSM8 A*02:01 conserves the extensive hydrogen bond and hydrophobic contact network to the C- and N-termini of the TAX9 peptide (fig. 4B).

**Figure 4.**
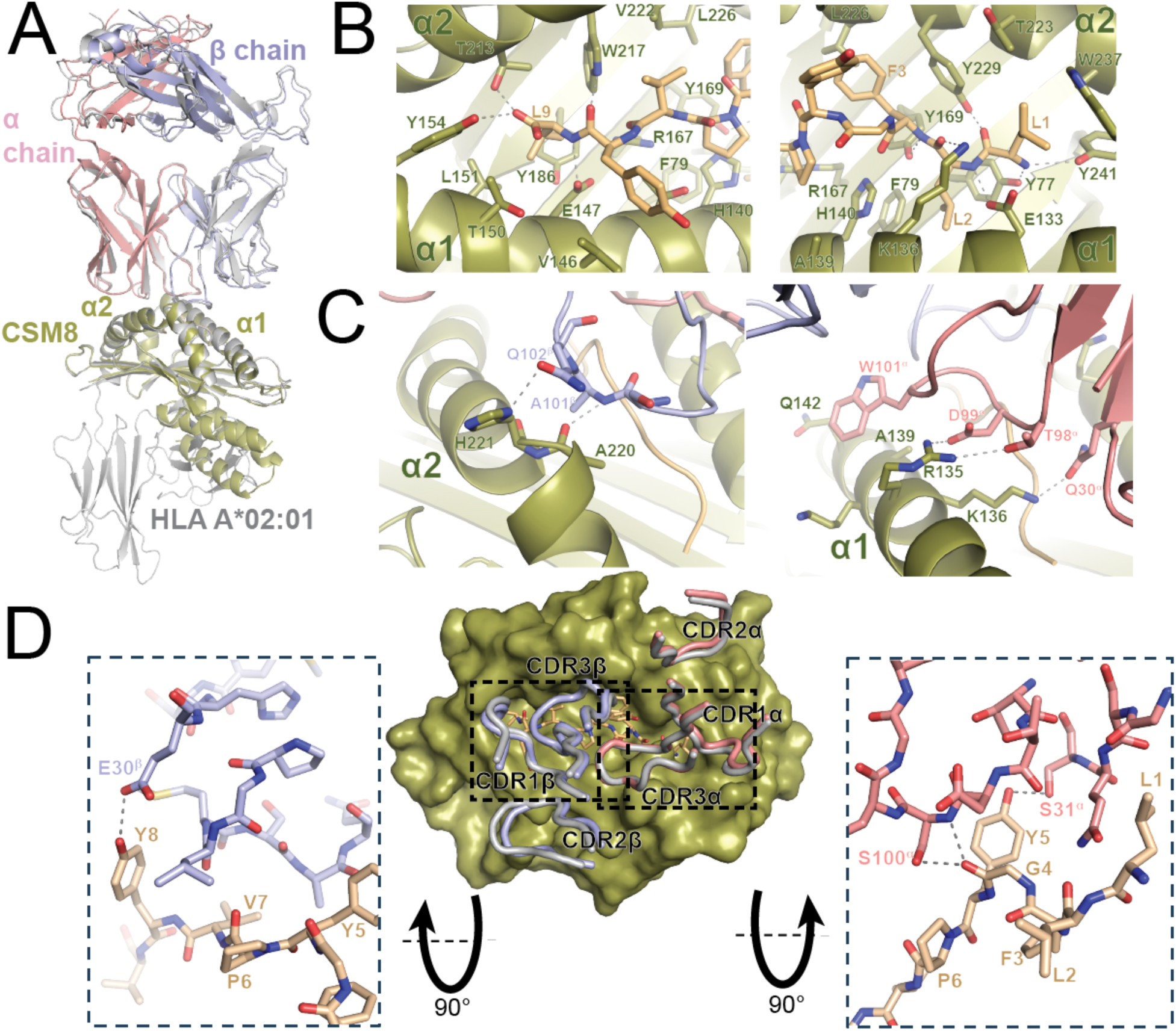
The crystal structure of the CSM8 A*02:01/TAX9/A6c234 complex reveals a native-like TCR docking footprint. A) crystal structure aligned to the HLA chain of the native A*02:01-TAX/A6c134 structure (PDB ID 4FTV, gray). For the current structure, the α chain is shown in salmon, the β chain in light blue, TAX9 peptide in light orange, and CSM8 A*02:01 in light green. The α1 and α2 helices of the CSM8 molecule are labeled for reference. B) CSM8 A*02:01 interactions with the N-terminus (left) and C-terminus (right) of TAX peptide. Interacting residues are shown as sticks, and hydrogen bonds are shown as dashed lines. C) CSM8 A*02:01 interactions with the β (left) and α (right) chains of the A6c234 TCR. D) Top view of CSM8 A*02:01 (shown as surface representation) with TCR CDR loops shown as tubes (center); CDR loops from the native structure are shown in gray. Peptide contacts from the β chain (left) and α chain (right) are shown as insets.

We further found that the TCR variable domains engage CSM8 A*02:01 nearly identically to native A*02:01 (Cα-RMSD 0.54 Å for 69 atoms in the complementarity determining region (CDR) loops). Though not all side chains are clearly visible in the electron density, those that are observed suggest that most or all residue contacts from the native complex are retained. In particular, we observe hydrogen bonds from the TCR β chain to the α2 helix of CSM8 A*02:01 including Glu102^β^ to H221 and Ala101^β^ to Ala220. At the α2 helix, the TCR α chain contributes hydrogen bonds from Asp99^α^ and Thr98^α^ to Arg135 and Gln30^α^ to K136 and also an extensive hydrophobic interaction between Trp101^α^ and the surface formed by Gln132, Ala139, and Lys138 (fig. 4C). A6c134 also makes four hydrogen bonds to TAX9: Glu30^β^ to Tyr8, Ser100^α^ to Gly4, Ser31^α^ to Tyr5, and a mainchain-mainchain contact between Ser100^α^ and Gly4 (fig. 4D). The close agreement of the CSM8 and native A*02:01/TAX9/A6c134, in conjunction with peptide and TCR binding data, suggests that, despite some issues with soluble expression,SMART A*02:01 can present peptides in the same conformation as native A*02:01.

### The designed stabilizing domain solubilizes several additional common HLA allomorphs

We expressed 15 human HLA allomorphs using our SMART stabilizing domain in *E. coli* and carried out small-scale purification using HPLC. SMART versions of many allomorphs exhibited peaks in the expected monomeric range, though many also displayed dimeric and aggregate peaks (fig. 5A). Interestingly, HLA A*03:01 and HLA A*01:01 demonstrated high protein expression, and HLA B*07:02 showed the highest monomeric fraction (fig. 5B). The presence of a significant dimeric or aggregated population, particularly in allomorphs with high expression, indicates that the stabilizer was not effective in simultaneously promoting high protein expression and monomeric behavior across diverse HLAs. However, we observed higher expression levels in four of the 15 allomorphs, and reduced dimerization in three, when compared to SMART A*02:01. These results suggest that the stability and expression of these allomorphs could be improved through minimal redesign of the stabilizing domain.

**Figure 5.**
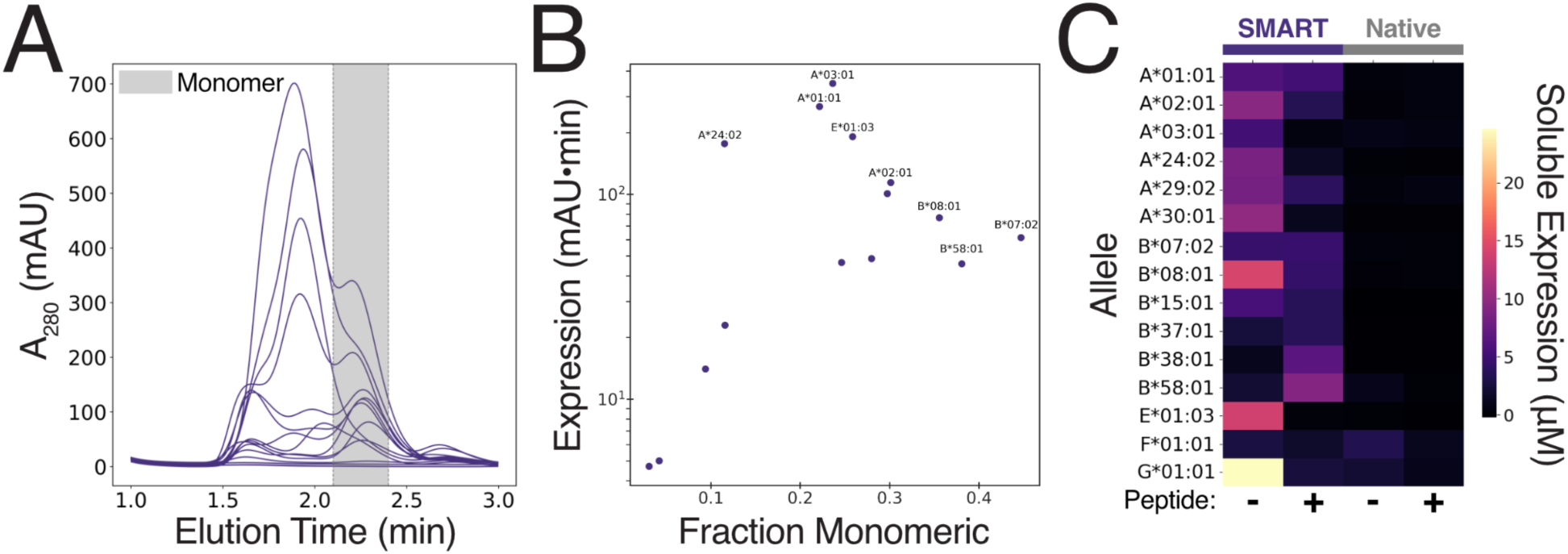
Stabilization of multiple HLA alleles using the SMART system. A) SEC traces of empty HLA alleles fused to SMART stabilizer. Expected monomer range for the S75 column on the HPLC is 2.1-2.4 minutes. B) Plot of SMART HLA expression statistics showing total area under the SEC curve versus the fraction of the sample that is monomeric. C) Heatmap of native and SMART HLA soluble expression levels with (+) and without (-) a linked peptide expressed *in vitro*. Soluble protein (µM) was quantified using radioactive ^14^C-leucine incorporation. Average of three replicates (n = 3) is shown for each construct.

We next compared the expression of native, full-length HLAs (native) with their SMART versions, with and without genetically linked peptides, using CFE. To determine whether SMART HLAs have improved solubility over native HLAs, we measured soluble protein using radiolabeled ^14^C-Leucine incorporation which allows for precise quantification of protein yields (fig. 5C). We found that almost all SMART HLAs had improved soluble yields compared to native HLAs, and that many SMART HLAs that did not express well in *E. coli* had significant CFE expression levels. The improvement in soluble yields in the CFE system could arise from reduced crowding under more dilute conditions, a more oxidizing redox state, or the presence of the DsbC chaperone. Both *E. coli* and CFE experiments suggest that the designed stabilizing domain enables the solubilization of different allomorphs without the need for extensive inclusion body preparation. Further design optimization could further improve expression and reduce aggregation.

### Peptide-fused SMART MHC oligomers stain T-cells in a TCR-specific manner

An important application of pMHCs in immunological research is the staining and identification of T-cells using pMHC tetramers^1^. We therefore tested whether SMART MHCs could be converted into a similar multimeric staining reagent and used for similar purposes. Rather than using streptavidin to tetramerize SMART MHCs, as is typically done, we chose to directly fuse them to a *de-novo* designed oligomeric protein which assembles into a tetrahedral architecture containing 12 subunits^57^. This allowed us to skip the biotinylation step which is necessary for streptavidin-based tetramerization^58^. To improve folding and assembly of oligomeric SMART MHCs, we fused a peptide of interest to the N-terminus of the SMART MHC along with a SUMO tag. Co-expression of the Ulp1 protease allows the N-terminus of the peptide to be cleanly cleaved, allowing it to bind properly in the peptide binding groove. The Y84A mutation in the MHC sequence was used to allow the peptide linker to exit the peptide binding groove^59^, and on the C-terminus of the oligomer, we fused a Myc tag to allow for antibody staining (fig. 6A-C).

**Figure 6.**
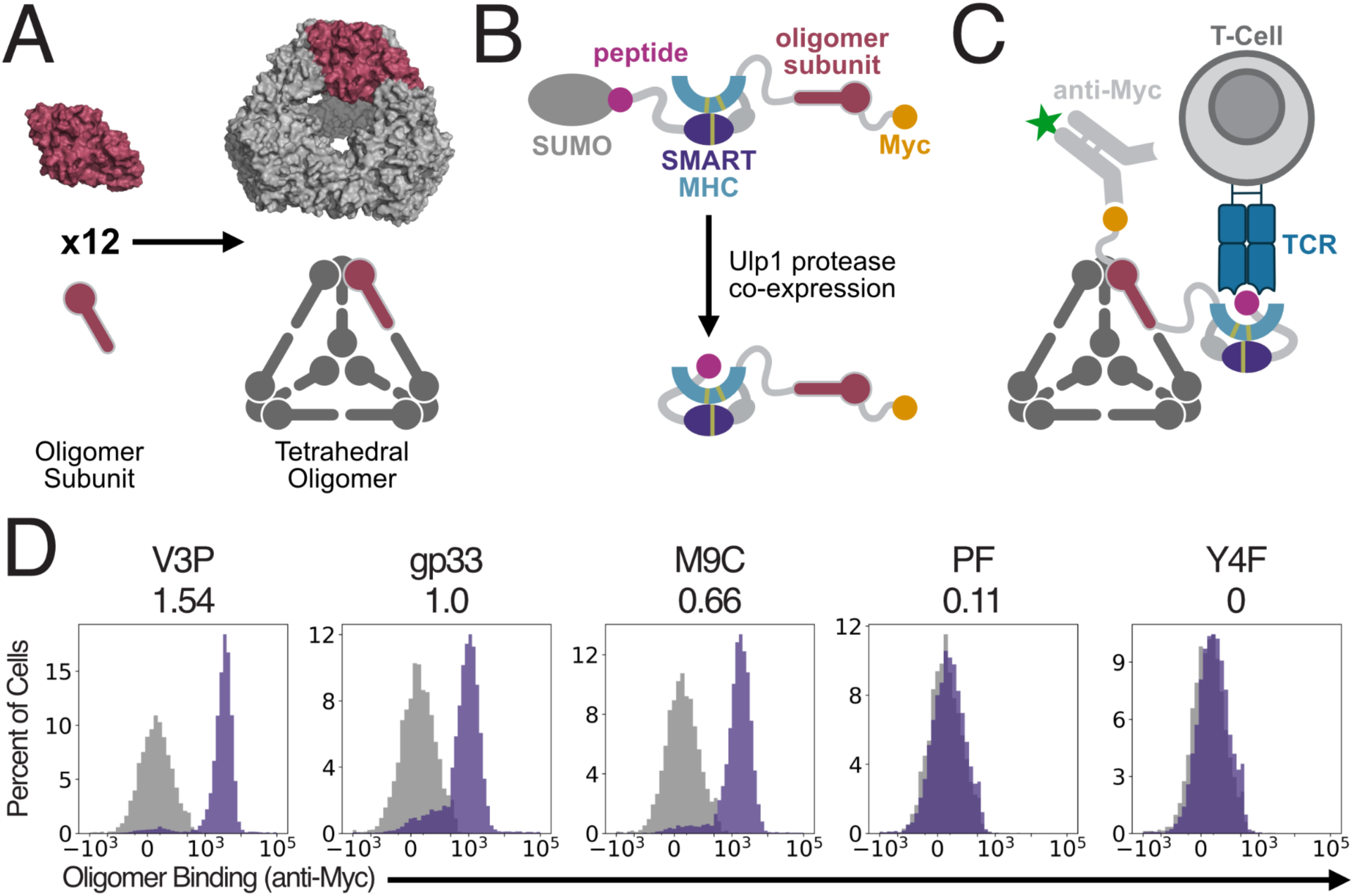
Peptide-fused SMART MHC oligomers stain T-cells in a TCR-specific manner. A) Twelve identical subunits assemble to form a tetrahedral oligomer. A single subunit is highlighted in red. B) Schematic of the SMART MHC oligomer sequence. Ulp1 protease is used to cleave the SUMO tag from the peptide, leaving a clean peptide N-terminus to bind the SMART MHC peptide binding groove. The MHC sequence bears the Y84A mutation to allow the C-terminal linker from the peptide to leave the binding groove. The peptide-fused SMART MHC is then fused to an oligomerization domain, followed by a Myc tag. C) The SMART MHC oligomer self-assembles into the same shape as the base oligomer in (A). Each subunit presents a peptide-fused SMART MHC to the T-cell and a Myc tag for secondary staining, but only one subunit is shown with these for simplicity. D) Flow cytometry measurements of T-cell staining of P14 T-cells (purple) or control TCR T-cells (gray) with SMART H-2D^b^ oligomers fused to several variants of the gp33 peptide. Numerical values under the peptide names indicate the binding strength of each peptide in the native MHC context for the P14 TCR relative to gp33 (K_D,gp33_/K_D,mut_). Values for V3P, PF, and Y4F were previously published by Duru *et al.*^49^. Value for M9C was published previously by Boulter *et al.*^30^

To assess the ability of the SMART MHC oligomers to stain T-cells, we mixed two populations of T-cells: one expressing the P14 TCR and the other expressing the unrelated TCR, MAG-IC3^60^, along with mTagBFP to allow the two cell types to be distinguished independently of TCR staining. We then expressed SMART H-2D^b^ oligomers fused to gp33 peptide variants with known affinities to the P14 TCR^30,49^ and assessed their ability to stain the T-cell mixture. We found that broad trends in TCR/pMHC binding affinity were reflected in our T-cell staining data; the highest affinity variant (V3P) clearly showed the brightest staining, with intermediate affinity peptides (gp33 and M9C) showing somewhat reduced staining, and low affinity variants (PF and Y4F) showing no staining. However, there were some deviations from our expectations; the M9C variant produced brighter staining than gp33 despite having a weaker affinity^30^, and the PF variant produced no staining despite its known (albeit weak) interaction with the P14 TCR^49^. These differences may result from changes in peptide conformation induced by the linker to the peptide, or by minor structural differences between SMART and native H-2D^b^. Nevertheless, staining was both TCR- and peptide-specific, suggesting that SMART MHC oligomers could be used alongside pMHC tetramers to more easily identify and isolate pMHC-specific T-cells.

## Discussion

Our success in creating a single stabilizing domain that allows soluble expression in *E. coli* of both the H-2D^b^ and A*02:01 MHC allomorphs paves the way for a generalizable stabilizing scaffold for other MHC-I allomorphs. Our structural and biochemical data confirm that the SMART versions of these two alleles maintain critical peptide and TCR interactions. Our structural data show that, for both allomorphs, the conformation of the presented peptide is nearly identical to the native conformation, and that for A*02:01, the TCR binds through the same set of interactions in both the native and SMART complexes. Thus, SMART MHCs can provide a convenient and shelf-stable complement to conventional H-2D^b^ and A*02:01, largely bypassing the need for refolding, and allowing more flexibility in the production of complexes with varied peptides. Although the designed stabilizing domain does not in its present form rescue all HLA alleles, it does improve biochemical properties of several additional allomorphs, suggesting that variants of the stabilizing domain could be developed to fully solubilize these and other HLA alleles.

In addition to their utility in biophysical and T-cell staining experiments, SMART MHCs could improve high-throughput measurements of pMHC/TCR interactions. Their high expression levels in yeast display could increase sensitivity of these experiments, potentially allowing detection of weaker interactions. Additionally, the soluble expression of SMART MHCs in cell-free systems opens many possible routes to characterize many pMHC-TCR interactions rapidly by taking advantage of the high-throughput and scalable nature of CFE^37,41,61,62^. Finally, the ability to produce a tetramer-like T-cell staining reagent without the need for more labor-intensive refolding protocols has the potential to reduce the barriers to performing T-cell tracking or sorting experiments. The SMART system could dramatically increase the number of peptide and MHC variants that can be tested, enabling improved tracking of immune responses to infectious disease, identification of cancer-targeting T-cell clones, and many other applications. To fully realize this potential, the SMART design will need further improvement, as some human alleles are not expressible in functional form with the SMART domain fusion (fig. 5). With such improvements, SMART MHCs have the potential to rapidly accelerate our understanding of T-cell behavior and TCR specificity.

## Methods

### Stabilizer library design

We used previously developed computational methods to design protein binders for arbitrary target proteins^28^ to design this stabilizing domain. The “target” supplied to this method was the α1 and α2 domains of the H-2D^b^ structure (PDB: 1S7U). We ran two versions of the design protocol. The first version was a completely *de-novo* approach which allowed any amino acid to be placed anywhere near the bottom of the H-2D^b^ peptide binding groove in order to make a favorable interaction. The second approach specifically focused on making designs which placed amino acids in locations that allow them to replace the exact interactions that β2m makes in the native structure. Designs from both of these protocols were pooled and filtered for the quality of the stabilizer and the interactions it made with the MHC, as previously described^28^. We also included a set of negative control designs which were made by randomly scrambling the sequence (while conserving the pattern of hydrophobic and hydrophilic residues) of a random subset of the designs.

### Yeast display screening of stabilizing domains

Yeast display was performed as previously described^28^ with the following changes. Rather than screening our designs for binding to the α1 and α2 domains of H-2D^b^, we fused the designs to those domains using a flexible poly-GS linker and screened them for surface display and binding to a FITC-labeled gp33 peptide (KAVYNFATM), with FITC linked to the amine on the lysine sidechain). Sorted populations were subsequently cultured and plasmid DNA was extracted for sequencing. FITC-gp33 was purchased from GenScript as a custom peptide synthesis.

### Protein expression and purification

Genes encoding the designed protein sequences were synthesized and cloned into modified pET-29b(+) E. coli plasmid expression vectors with a 6xHis tag added to the N-terminus (for monomeric versions) or the C-terminus (for oligomeric/peptide-fused versions). Plasmids were transformed into chemically competent E. coli BL21 (DE3) cells (NEB). E. coli cells were grown in LB medium at 37 °C until the cell density reached 1.0 at OD600. Then, IPTG was added to a final concentration of 1mM and the cells were grown overnight at 16°C for expression. The cells were collected by spinning at 4,000 g for 5 min and then resuspended in lysis buffer (150 mM NaCl, 25 mM Tris-HCL (pH 8.0), 25 mM imidazole, 1mM PMSF, and 5% glycerol) with RNase. The cells were lysed with a Qsonica Sonicators sonicator for 7 min in total (3.5 min each time, 10 s on, 10 s off) with an amplitude of 80%. The soluble fraction was clarified by centrifugation at 14,000 g for 30 min. The soluble fraction was purified by immobilized metal affinity chromatography (Qiagen) followed by FPLC SEC on a Superdex 75 10/300 GL column (GE Healthcare) for monomeric versions, and a Superose 6 10/300 GL column (GE Healthcare) for oligomeric versions. All protein samples were characterized by SDS–PAGE, and purity was greater than 95%. Protein concentrations were determined by absorbance at 280 nm measured with a NanoDrop spectrophotometer (Thermo Scientific) using predicted extinction coefficients.

### Cell-free gene expression (CFE) and radioactive quantification of soluble protein yields

Fifteen µL CFE reactions were prepared in 2 mL microcentrifuge tubes (Axygen) using an adapted version of the existing PANOx-SP formulation to create an oxidizing environment^38–40^. The reaction environment included: 8 mM magnesium glutamate, 10 mM ammonium glutamate, 130 mM potassium glutamate, 1.2 mM ATP, 0.85 mM GTP, 0.85 mM UTP, 0.5 mM CTP, 30 µg/mL folinic acid, 0.17 mg/mL *E. coli* tRNA, 0.40 mM NAD, 0.27 mM CoA, 4.00 mM oxalic acid, 1.00 mM putrescine, 1.5 mM spermidine, 57 mM HEPEs, 2 mM 20 amino acids, 0.03 M phosphoenolpyruvate, 4 mM oxidized glutathione, 1 mM reduced glutathione, 10 µM purified *E. coli* DsbC, 30% v/v *E. coli* extract treated for 30 minutes with 50 µM iodoacetamide at room temperature, and nuclease free water. All components were added together on ice^41,43^. 10 µM radiolabeled leucine, ^14^C-leucine (Revvity NEC279E250UC, 11.1GBq/mMole), was added to the final reaction mixture. This final reaction mixture was added to 6.66% v/v unpurified linear DNA PCR products (LETs) in triplicate 15 µL reactions^62,63^. Tubes were incubated at 30°C overnight. Samples were spun at 16,000xg for 10 minutes at 4°C to separate total and soluble protein fractions. 6 µL of supernatant were incubated at 37°C for 20 minutes with 0.25 N KOH. Fractions were spotted on a 96 well filtermat (Revvity 1450–421), dried, and precipitated by 5% trichloroacetic acid at 4°C (washed). Filtermats were dried and scintillation wax was melted onto the mat. A scintillation counter (Revvity MicoBeta2) measured radioactivity.

### Mass Spectrometry

To identify the molecular mass of each protein, intact mass spectra was obtained via reverse-phase LC/MS on an Agilent G6230B TOF on an AdvanceBio RP-Desalting column, and subsequently deconvoluted by way of Bioconfirm using a total entropy algorithm.

### Peptide synthesis

Fmoc protected amino acids were purchased from P3 Biosystems. OxymaPure and preloaded Wang resins were purchased from CEM. DIC was purchased from Oakwood Chemicals. HATU, N,N-diisopropylethylamine (DIEA), and tris(2-carboxyethyl)phosphine (TCEP), Fmoc-Lys(MTT)-OH, trifluoroacetic acid (TFA), triisopropylsilane (TIPS), 3,6-dioxa-2,8-octanedithiol (DODT) were purchased from Sigma-Aldrich. All solvents were purchased from Fisher Scientific. AF488 carboxylic acid was purchased from Lumiprobe. Unmodified peptides were synthesized on preloaded Wang resin on a CEM LibertyBlue microwave synthesizer with standard solid-phase peptide synthesis protocols, transferred to a reaction vessel, and treated with a cleavage solution of 92.5:2.5:2.5:2.5 TFA:H2O:TIPS:DODT for 3h to obtain the crude peptide. After the cleavage reaction, the cleavage solution was precipitated into ice cold ether, spun down, washed with ether and dried under nitrogen to obtain a crude peptide pellet. The crude peptide was then purified on an Agilent 1260 Infinity semi-preparative RP-HPLC with a linear gradient of A (H2O with 0.1% TFA) to B (acetonitrile (ACN) with 0.1% TFA). Fractions were isolated and checked for the correct mass using an Agilent G6230B TOF, then combined and lyophilized to dryness to obtain pure peptide.

### Fluorophore-conjugated peptide synthesis

Synthesis proceeded as described above, substituting the Fmoc-Lys(MTT)-OH for the desired modified Lys in the sequence. The synthesized peptide was then transferred to a reaction vessel and washed with DCM, then treated with a solution of 2% TFA and 2% TIPS in DCM to orthogonally deprotect the MTT group. The resin was washed with DCM and DMF, then treated with the AF488 carboxylic acid (3eq) in a coupling solution of HATU (3eq), DIEA (5eq), in DMF, to react protected from light for 3h. After completion of the reaction, the resin was washed with DMF and DCM, then proceeded through the cleavage and purification protocol as described above.

### Cleavage site mutation design

Cleavage site design was used to make mutations to the MHC sequence that would reduce proteolysis without impacting peptide or TCR binding. First, a multiple sequence alignment of MHC protein sequences was collected using PSI-BLAST^64^. Next, the MSA was converted into a position-specific score matrix (PSSM) denoting the likelihood of observing each possible amino acid at each position in the MSA. Standard Rosetta design protocols^65^ were modified to allow mutations to amino acids that had likelihoods above a specified cutoff, and to only allow mutations at sequence positions near the pre-identified cleavage. Design was further restricted to prevent mutations in residues with sidechains that could interact with a bound peptide or TCR. All cleavage site mutants were designed based on the hit6 design model as a starting point, and the highest scoring designs were selected for experimental testing based on a combination of Rosetta metrics relating to the overall Rosetta energy of the design model, the complementarity of the stabilizer/MHC interface, and the number hydrogen bond donors/acceptors that were left unbonded.

### Linker design

Linker design was used to replace the flexible GS linker that was used in the yeast display screening with a more structured and shorter linker. Using the design model of the CSM8 variant as a starting point, we used previously developed “inpainting” methods^34^ to fill in a small segment of protein structure to bridge the gap between the C-terminus of the stabilizer and N-terminus of the MHC. Sequences for the protein backbones produced by this method were designed using ProteinMPNN^35^, restricted to changing only the amino acids in the “inpainted” structure. Finally, the resulting designs were evaluated using AlphaFold^36^ predictions where the MHC structure was provided as a template, but the stabilizer and linker were not. Designs with high overall pLDDT scores and low PAE scores for residues in the MHC/stabilizer interface were selected for experimental testing.

### Peptide binding affinity measurements

Three technical replicates of varying concentrations of SMART MHC were mixed with a constant concentration of fluorophore-labeled peptide (300pM FITC-gp33 for H-2D^b^ and 10nM AF488-NY-ESO-1 for A*02:01) and incubated overnight at room temperature to allow equilibration. Fluorescence polarization measurements of these samples were made with a Synergy Neo2 plate reader (BioTek instruments) with a 485/530 FP filter. Binding curves (using the non-simplified equilibrium binding equation^66^) were fitted separately to each of the triplicate measurements and averaged to determine the K_D_. The peptides used in these experiments were synthesized in-house using the methods described above.

### Circular dichroism measurements

Purified soluble SMART H-2D^b^ was diluted to 0.3mg/mL (9.0μM) in 25mM Tris (pH 8.0), 150mM NaCl, and 5% glycerol. The gp33 peptide stock was prepared by dissolving dry peptide (purchased from GenScript as a custom synthesis) to 2mg/mL (1.52mM) in methanol. Two-fold molar excess of peptide stock (or an equivalent volume of methanol) was added to the protein sample and incubated at 4°C overnight. CD spectra were measured using a Jasco J-1500, and melting curves were measured in increments of 0.5°C at a rate of 2°C/min.

### Surface Plasmon Resonance (SPR) measurements

TCR binding affinities were measured as previously described^49^, using CSM8 H-2D^b^ as the mobile phase instead of native H-2D^b^. All measurements were performed on a BIAcore T200 (GE Healthcare) at 20°C in the buffer containing 10 mM HEPES pH7.4, 150 mM NaCl, 0.005% Tween-20, 3 mM EDTA. Soluble P14-his6 was noncovalently coupled to the anti-his antibody, immobilized on a CM5-chip via standard amine coupling, and around 4000 response units of anti-his antibody was coupled, immobilizing around 1000 response units of P14-his6 (0.75 µM). A control surface was generated the same way, and up to 100 μM or 200 μM of freshly produced CSM8 H-2D^b^/peptide complexes (at least ten 2-fold dilutions from stock, three of which were repeated in duplicate) were injected over the chip surfaces at 30 μL/min. The sample rack was cooled to 4°C during the run. Chip surfaces were regenerated using 0.1 M Glycine-HCl pH 2.5, 500 mM NaCl, Tween 0.05% at 30 μL/min after each injection. The CSM8 H-2D^b^/peptide was injected over a control surface, and the final signal was calculated by subtracting the signal obtained on the control surface from the signal on the TCR-coupling surface, to remove the contributions of the bulk effect and possible non-specific binding. K_D_ values were calculated by fitting a standard binding curve with unknown maximum binding and K_D_ to the equilibrium binding values determined by BIAevaluation 3.0 software.

### CSM8 H-2D^b^ Crystallography

Crystallization of CSM8 H-2D^b^/gp33 was performed using the sitting-drop vapor diffusion method at 293.15 K. Drops were set up using 0.15 µL of the SMART H-2D^b^/gp33 and 0.15 µL of the reservoir solution (0.2 M sodium acetate trihydrate, 0.1 M TRIS hydrochloride pH 8.5, and 30% w/v polyethylene glycol, PEG 4,000), equilibrated against 50 µL of reservoir solution. Crystals appeared after 6∼13 days and were cryoprotected with a solution containing an additional 7.5% w/v PEG 4,000 in the reservoir solution and harvested using mounted CryoLoops (Hampton Research). Subsequently, the crystals were flash-frozen in liquid nitrogen and transported to the beamline for data collection. Diffraction images were collected in the automatic beamline ID30 at the European Synchrotron Radiation Facility (ESRF) in Grenoble, France. Diffraction data were processed using autoPROC^67^. The crystal structure was determined by molecular replacement utilizing Phaser-MR in PHENIX^68^ with the design model employed as the search model. Subsequent refinement was carried out using PHENIX. Manual model building was conducted using Coot^69^, and the resulting model was further refined using PHENIX^70–72^. The figures were generated using PyMOL Molecular Graphics System (Schrödinger). The final coordinates/structure factors have been deposited in the PDB with accession code 9HY4.

### Peptide-fused yeast display

Yeast display of peptide-fused SMART A*02:01 was performed as previously described^19^. In brief, 50 ng pCT or pYAL plasmids encoding corresponding full length or SMART A02 constructs with TAX were electroporated into competent EBY100. The EBY100 was cultured in YPD medium for 1 hour at 30°C, spun down and continued to grow in SDCAA medium for 48 hours before induction in SGCAA for 48 hours. Display levels were evaluated with fluorophore-conjugated anti-HA and anti-A*02:01 (clone BB7.2) antibodies, and TCR binding was evaluated with TCR tetramers made by combining soluble biotinylated A6 TCR with fluorophore-conjugated streptavidin.

### A6c134 TCR Purification

The A6c134 protein was produced using the Expi293F™ GnTI-Cells (Thermo Fisher Scientific). In brief, the α chain of A6c134 was cloned into pD649 expression vector with an acidic zipper and 6x Histag, the β chain was cloned with a basic zipper and 6x His tag. 1 μg TCRα and 1 μg TCRβ construct were transfected into 106 Expi293 GnTI-cells according to the manufacturer’s protocol. The supernatant of the transfected Expi293 GnTI-cells was harvested 4 days later, diluted with equal volume of PBS and 20mM (final concentration) Tris.Cl (pH 8.0). The supernatant was incubated with 2ml Ni-NTA (Thermo Fisher Scientific) at 4oC overnight, the Ni-NTA was washed twice with 10 mM Imidazole and the bound protein was eluted using 300 mM Imidazole. The TCR protein was buffer-exchanged with HBS and concentrated with a 30 kDa filter (Millipore). The protein was then treated with carboxypeptidase A/B, Endo H and 3C protease to remove sugars and zippers at 4°C overnight. The protein was purified by size-exclusion chromatography using a Superdex S200 Increase column on AKTA Pure FPLC (Cytiva) and the size was confirmed using a SDS-PAGE gel.

### CSM8 A*02:01 expression and refolding

Codon-optimized DNA encoding SMART A*02:01 CSM8 L11 was expressed in BL21 (DE3) E. coli cells into inclusion bodies by induction with 1 mM IPTG at OD600 = 0.6 followed by cell incubation at 37°C for 5-6 hours at 200 RPM. After solubilizing the purified inclusion bodies in guanidine-HCl, ∼100 mg of protein were rapidly diluted dropwise into 1000 mL of chilled refolding buffer (Tris pH 8, 2 mM EDTA, 0.4M Arginine, 4.9mM L-glutathione reduced, 0.57 mM L-glutathione oxidized, and 2.5mg of TAX9 (LLFGYPVYV) peptide, solubilized in 50% DMSO) at 4 °C while stirring. Refolding proceeded 4 days at 4 °C without stirring. The solution was dialyzed into a buffer of 150 mM NaCl, 25 mM Tris (pH 8) overnight and concentrated using Tangential Flow Filtration. Purification of refolded SMART MHC was performed by SEC with a HiLoad Superdex 200 16/600 column (Cytiva) at 1 mL/min with running buffer (150 mM NaCl, 25 mM Tris (pH 8)), resulting in a final protein concentration of 25 mg/mL.

### A6c134/CSM8 A*02:01/TAX9 Crystallography

Purified A6c134 TCR was combined with equimolar refolded CSM8 A*02:01/TAX9 and purified on a Superdex S200 10/300 Increase column in HBS. Initial seed crystals were obtained with protein complex concentrated to 13 mg/mL and crystallized by vapor diffusion against a reservoir of 100 mM MES pH 6.5, 12% PEG 20,000, using the MCSG-2 crystallization screen (Anatrace) at 293 K. A second cycle of microseed matrix screening was performed using the Index screen (Hampton Research), with new seeds grown from 0.1 M Bis-tris pH 6.5, 25% PEG 3350, and final crystals were grown at 16 mg/mL using these seeds and a reservoir solution of 50 mM Mg(CHO)_2_, 17% PEG 3350. Crystals were cryoprotected with addition of 30% glycerol and flash frozen. Diffraction images were collected at NSLS-2 beamline 17-ID-1 at Brookhaven National Laboratory. Diffraction data were processed using XDS^73^ and aimless^74,75^. The structure was solved by molecular replacement using Phaser-MR in PHENIX, using the designed model of hit6 A*02:01 and PDB model 4GRM as search models to place two complexes in the asymmetric unit. Iterative cycles of rebuilding and refinement were performed using Coot and Phenix. Final refinement in Phenix at 3.4 Å resolution included individual B-factor and TLS refinement using automatically determined noncrystallographic symmetry (NCS) restraints, with torsional NCS sigma = 2.5° and limit = 10°. An example of electron density at the final stage of refinement is shown in Supplementary Figure S6. Data collection and refinement statistics are presented in Supplementary Table S2. Structure figures were generated using PyMOL. Structure factors and final model coordinates have been deposited in the PDB with accession code 9NDS, and diffraction images have been deposited in the SBGrid Databank.

### Small-scale expression and SEC of HLA Alleles

Linear DNA encoding native, full-length HLA alleles, with or without covalently linked peptides, were cloned into modified pET-29b(+) vector containing an N-terminal 6xHis tag via Golden Gate Assembly (BsaI-HF® v2). Plasmids were transformed into chemically competent *E. coli* BL21(DE3) cells (NEB) and received in LB media. Cells were diluted and grown in Terrific Broth II (TB-II) media + 50 µg/mL Kanamycin at 37°C in 96 well microplates with long drip spouts until reaching an OD600 of 1.0, followed by the addition of 1 mM IPTG. Plates were transferred to 16°C for protein expression. Plates were spun down for 5 minutes at 4000xg and cells were resuspended in a lysis buffer consisting of BugBuster® Protein Extraction Reagent, 0.1 mg/mL lysozyme, 0.01 mg/mL benzonase, 1 mM PMSF. Plates were shaken at 1000 rpm at 37°C for 15 minutes. Lysate was spun down and purified in 96 well 25 µm polyethylene fritted plates (Agilent) by immobilized metal affinity chromatography (Qiagen). Soluble, filtered samples were analyzed using high performance liquid chromatography (Agilent 1260 Infinity II LC System) on a Superdex™ 75 Increase 5/150 small-scale SEC column (Cytiva).

### T-cell staining

Two Jurkat T-cell lines were used: one expressing the P14 TCR, and the other expressing an unrelated TCR (a3a TCR, recognizing the MAGE-A3/A*01:01 complex) as well as mTagBFP. Both cell lines were mixed in equal numbers and resuspended to 1M cells/mL in 100uL of staining solution (500nM SMART MHC oligomer, 25mM Tris-HCl pH 8.0, 150mM NaCl, and 5% glycerol) and incubated at 4°C for 30min. Cells were then washed twice with 100uL of Fc block (HBH (HBSS with 0.5% BSA, and 10mM HEPES) supplemented with 10% 2.4G2 cell culture supernatant). Cells were then resuspended in 50uL AF647-anti-Myc antibody diluted in Fc block and incubated at 4C for 20min. Cells were then washed twice with 100uL of HBH and resuspended to a final volume of 150uL of HBH for analysis on an Attune Nxt flow cytometer. Cells were gated into P14 (mTagBFP-) or a3a (mTagBFP+) populations and the brightness of the AF647 stain for each population was compared.

## Data, Materials, and Software Availability

Atomic coordinates for the CSM8 H-2D^b^/gp33 and CSM8 A*02:01/TAX9/A6c13 will be uploaded to the PDB with the accession codes 9HY4 and 9NDS respectively. Code used for any computational design methods not published elsewhere will be uploaded to github. Sequences described in this paper are listed in supplemental tables S3-S6.

## Acknowledgements

We thank members of the Kueh lab for discussion and feedback on the manuscript, and Dr. Matthew Wither for providing the P14 Jurkat cell line. We also thank Dr. Brian Coventry and Dr. Lonxing Cao for the development of the protein design protocols used to create the yeast display library of stabilizing domains, and for their help adapting their protocols to the context of MHC stabilization. We also thank Dr. Sangmin Lee for the development of the oligomer used in our T-cell staining experiments. Finally, we thank Dr. Chris Norn and Kiera Sumida for the development of the protein design protocols used to create the CSMs, and for their help running those protocols. The Center for BioMolecular Structure (CBMS) is primarily supported by the National Institutes of Health, National Institute of General Medical Sciences (NIGMS) through Grant # P30GM133893, and by the DOE Office of Biological and Environmental Research FWP # BO070. This research used resources 17-ID-1 of the National Synchrotron Light Source II, a U.S. Department of Energy (DOE) Office of Science User Facility operated for the DOE Office of Science by Brookhaven National Laboratory under Contract No. DE-SC0012704. This work was funded by grants from the Bill and Melinda Gates Foundation (INV-010680; to David Baker), the National Institutes of Health (5 R01 AI103867 08; to K.Christopher Garcia), the Swedish Research Council (2021-05061; to Adnane Achour), the Swedish Cancer Society (24 3775 Pj O1 H; to Adnane Achour), the King Gustaf V Jubileum Fund (244092; to Adnane Achour), the Swedish Cancer and Allergy Foundation (11338; to Adnane Achour), the Defense Threat Reduction Agency (HDTRA1-21-1-0038; to Michael C. Jewett), the Army Research Office DURIP (W911NF-23-1-0334; to Michael C. Jewett), and a National Science Foundation GRFP award (DGE-1656518; to Laura Guerrero). This work was delivered as part of the NexTGen team supported by the Cancer Grand Challenges partnership funded by Cancer Research UK (CGCATF-2021/100002) and the National Cancer Institute (CA278687-01) and The Mark Foundation for Cancer Research. This work was delivered as part of the MATCHMAKERS team, supported by the Cancer Grand Challenges partnership financed by CRUK (CGCATF-2023/100006) and the National Cancer Institute (OT2CA297242). This work was delivered as part of the MATCHMAKERS team, supported by the Cancer Grand Challenges partnership financed by CRUK (CGCATF-2023/100008) and the National Cancer Institute (OT2CA297288). Portions of the text and figures were developed from the thesis of William L. White.

