## Supplemental Information for "Design of solubly expressed miniaturized SMART MHCs"

**Supplemental Figures and Information**

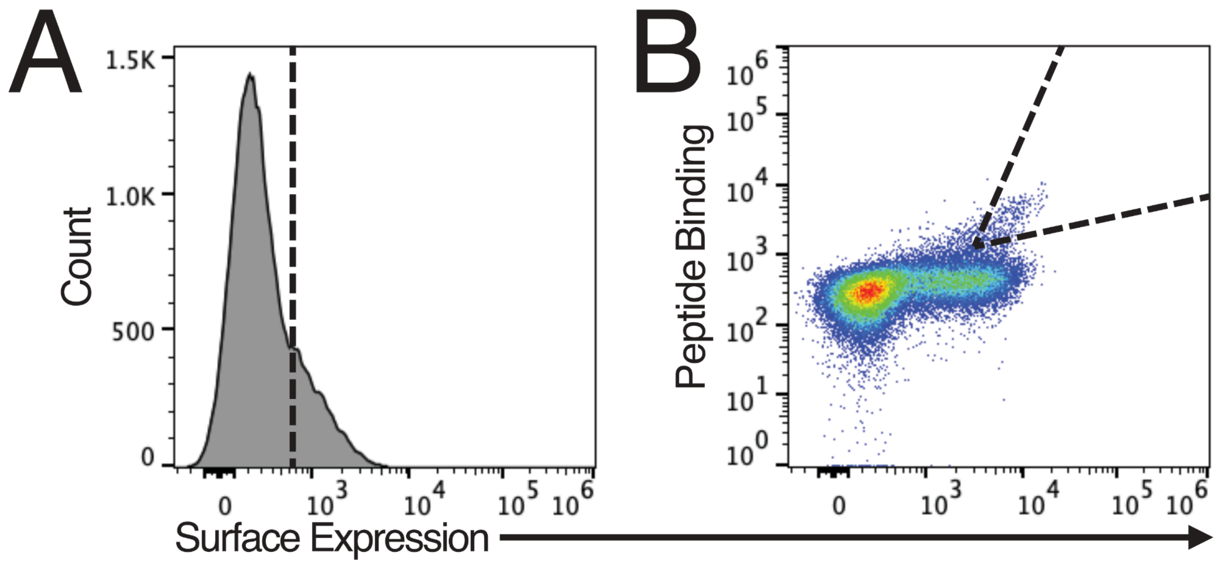

**Figure S1. Yeast display sorting of the stabilizer library.** A) Histogram of surface expression levels of the stabilizer library. Surface expression was evaluated by antibody staining of a C-terminal Myc tag. Cells with high surface expression were collected after the sort and recultured for the next sort. B) Joint distribution of the surface expression and peptide binding levels of the stabilizer library after the sort in A. Peptide binding was evaluated with FITC-labeled gp33 peptide. Cells with high surface expression and peptide binding were collected after the sort.

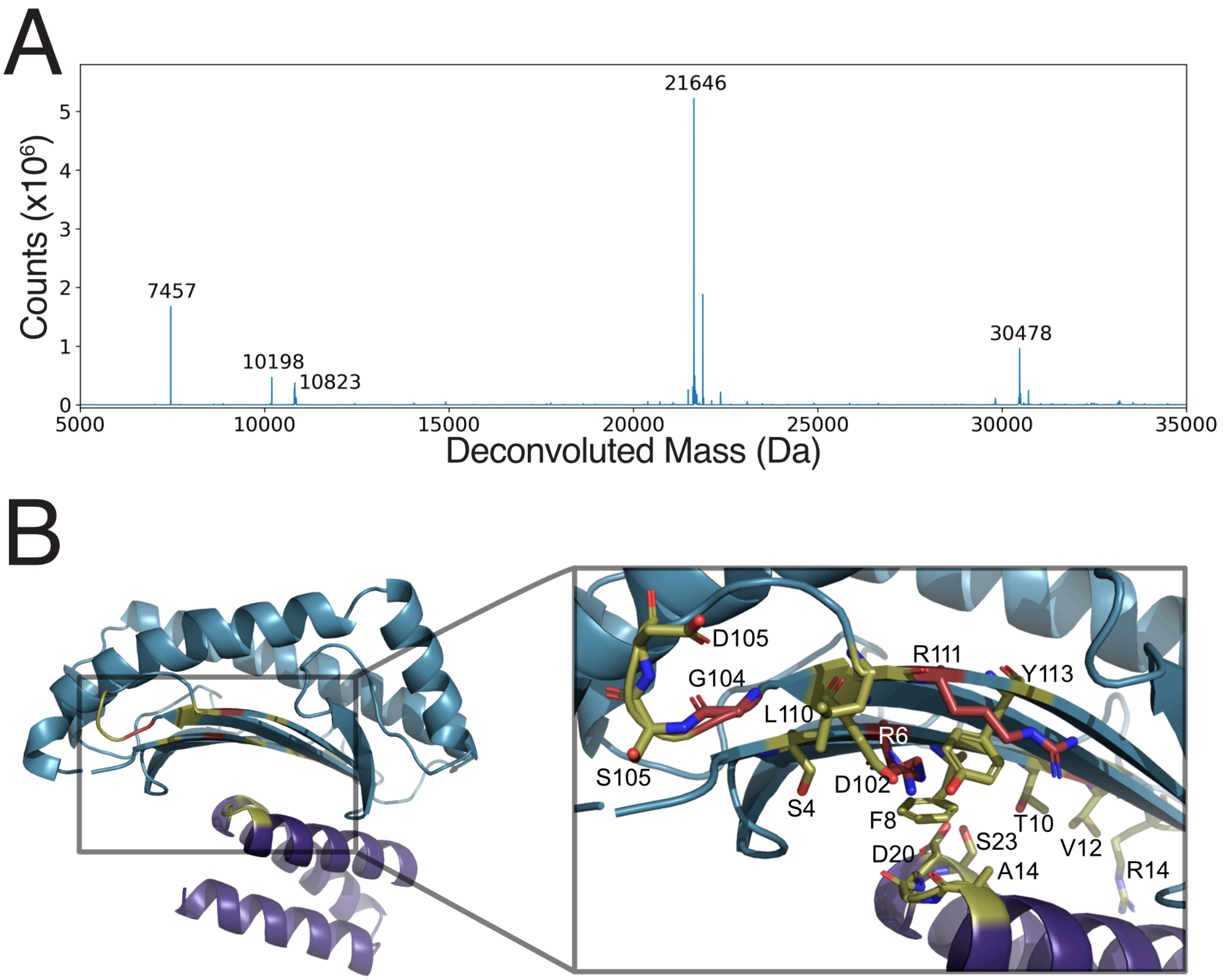

**Figure S2. MS analysis of hit6 cleavage sites.** A) Deconvoluted mass spectrum of the proteolytic fragments of hit6. Peaks that were used to identify cleavage sites are labeled with their mass in Da. B) Design model of hit6 with cleavage sites (red) and residues considered for redesign (sidechains showing; yellow) highlighted.

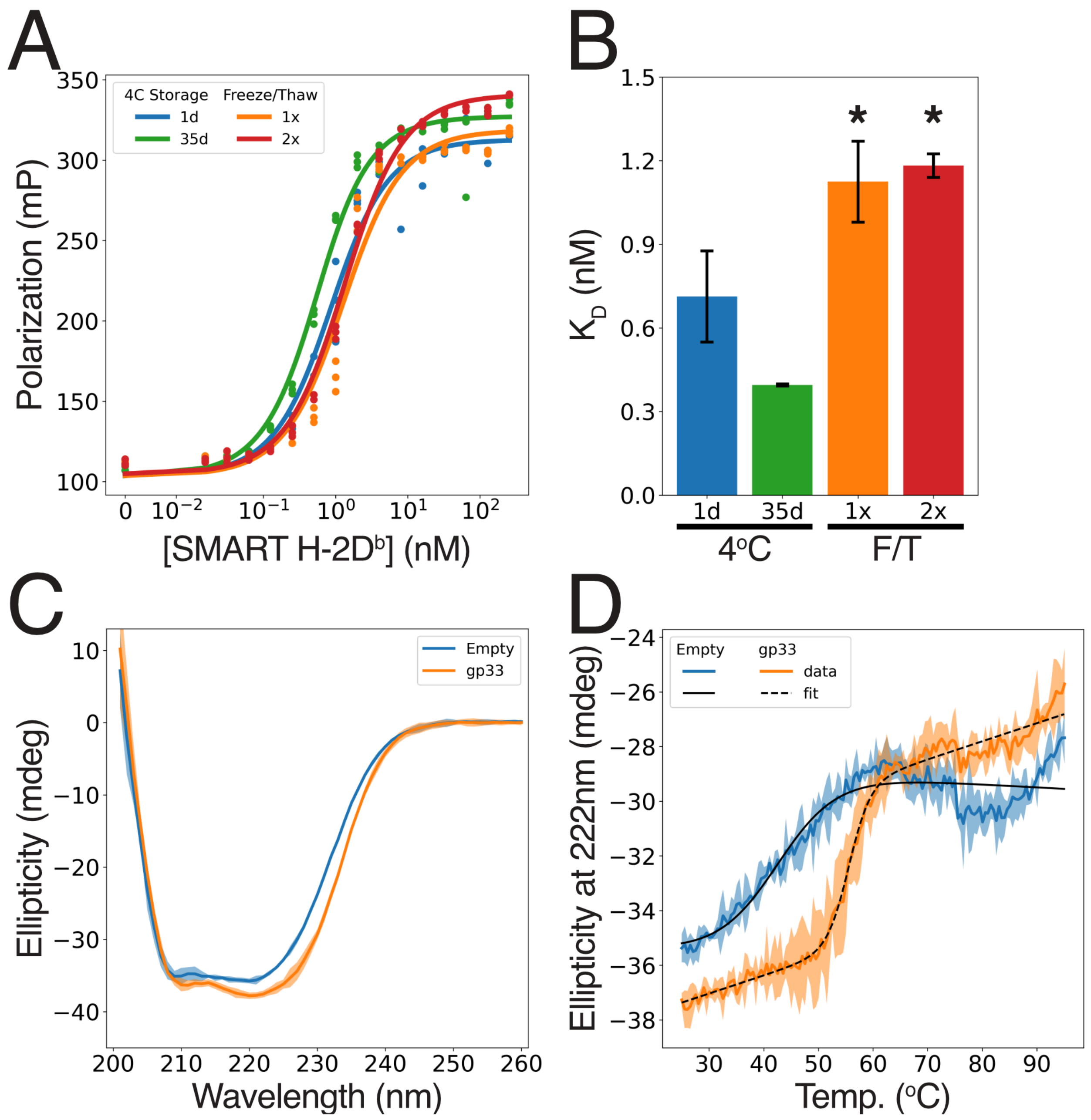

**Figure S3. Thermostability and shelf-life of SMART H-2D$^b$.** A) FP values (dots) and fitted binding curves (lines) for titrations of SMART H-2D$^b$ subjected to a variety of storage conditions. Data points represent each of n=3 replicates per condition. 1d or 35d: storage at 4°C for 1 or 35 days, respectively. 1x or 2x: flash frozen in liquid nitrogen and thawed at RT once or twice, respectively. FITC-gp33 peptide was used with all samples. B) Bar chart of the fitted $K_{D,app}$ of each sample. Error bars represent the standard deviation of 3 technical replicates. * indicates a p-value < 0.05 when compared to the 1d sample. C) CD spectra taken at 25°C comparing SMART H-2D$^b$ in the presence (orange) or absence (blue) of a 2-fold molar excess of the gp33 peptide. D) CD melting curves (colored lines) for the same samples as in (A). A sigmoid curve with an added linear component was fitted to each curve (black lines) to determine the melting temperature of each sample. In both (C) and (D), solid lines and shaded regions represent the mean and 99% confidence interval of 3 technical replicates, respectively.

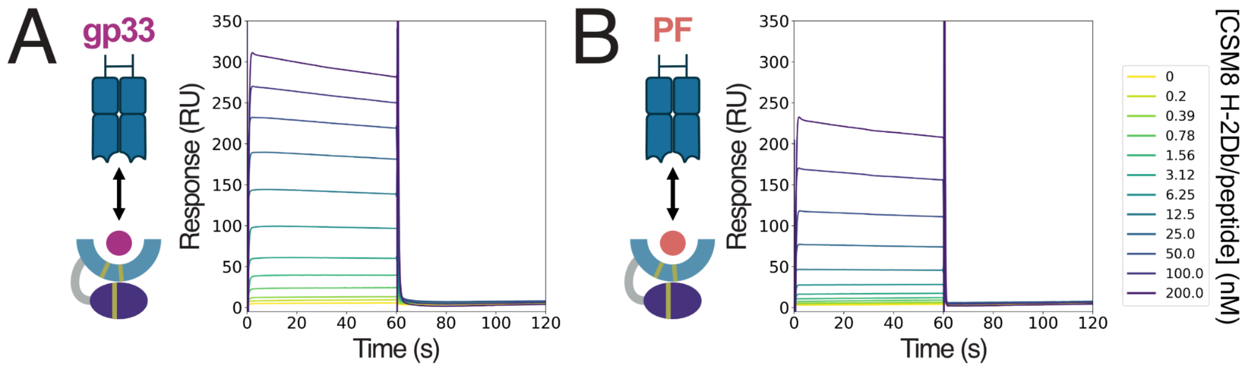

**Figure S4. P14 TCR binding kinetics.** Representative SPR traces from TCR pounding experiments with the P14 TCR and several variants of the gp33 peptide complexed to CSM8 H-2D$^{b}$. A) Data from the gp33 peptide. B) Data from the PF variant of the gp33 peptide.

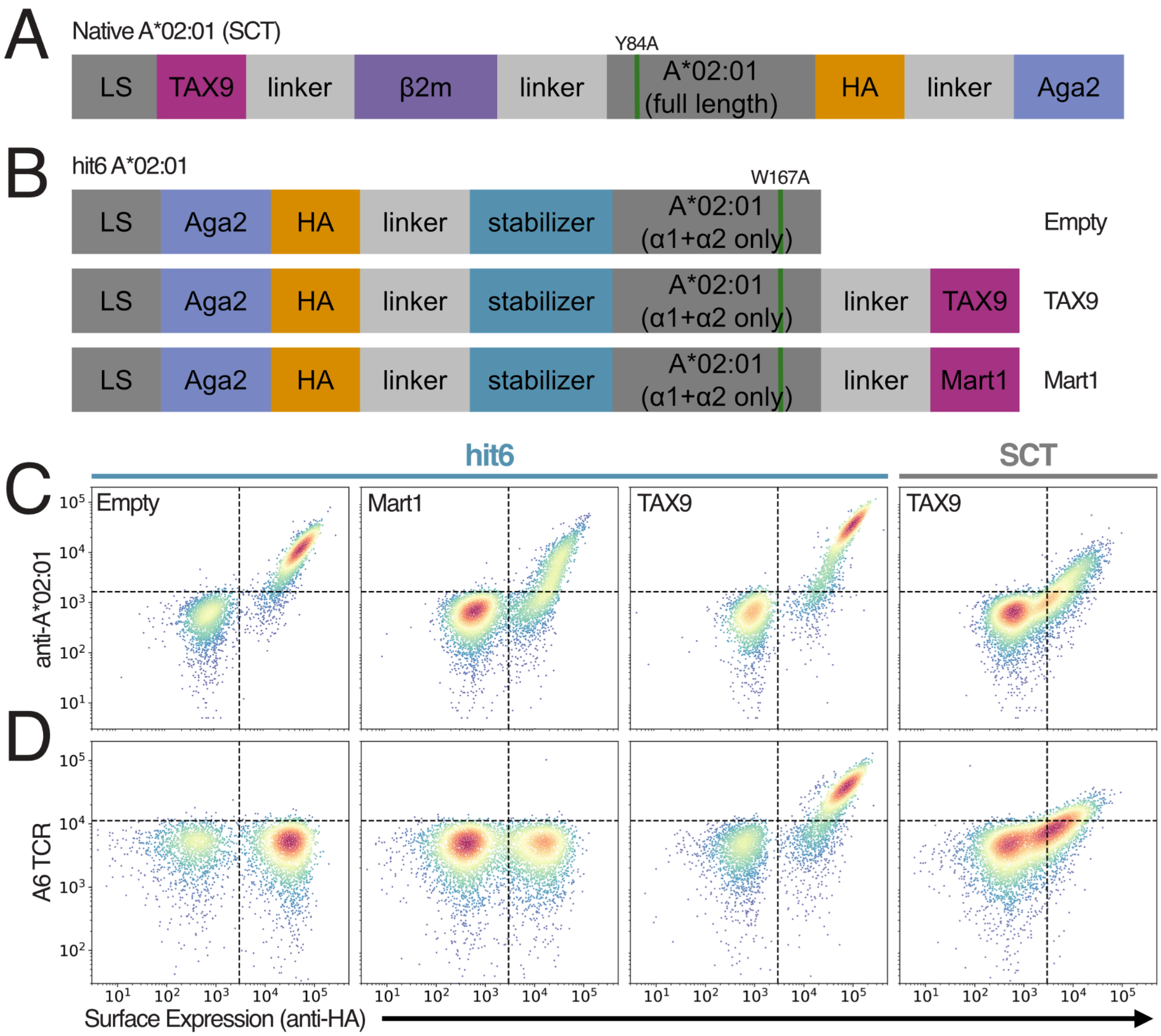

**Figure S5. Details of hit6 A*02:01 yeast display.** A,B) Schematics of native (A) and hit6 (B) A*02:01 yeast display constructs. LS: leader sequence; HA: hemagglutinin peptide tag, A) The Y84A mutation is used to allow room for the linker on the C-terminus of the peptide to leave the binding pocket. B) The W167A mutation is used to allow room for the linker on the N-terminus of the peptide to leave the binding pocket. C) Scatterplots from flow cytometry measurements of expression levels of various A*02:01 constructs on the surface of yeast. Surface expression was measured by binding of an anti-HA antibody, and by binding of an A*02:01-specific antibody. D) Scatterplots from flow cytometry measurements of A6 TCR binding of various A*02:01 constructs on the surface of yeast. TCR binding was measured with streptavidin tetramers of the A6 TCR.

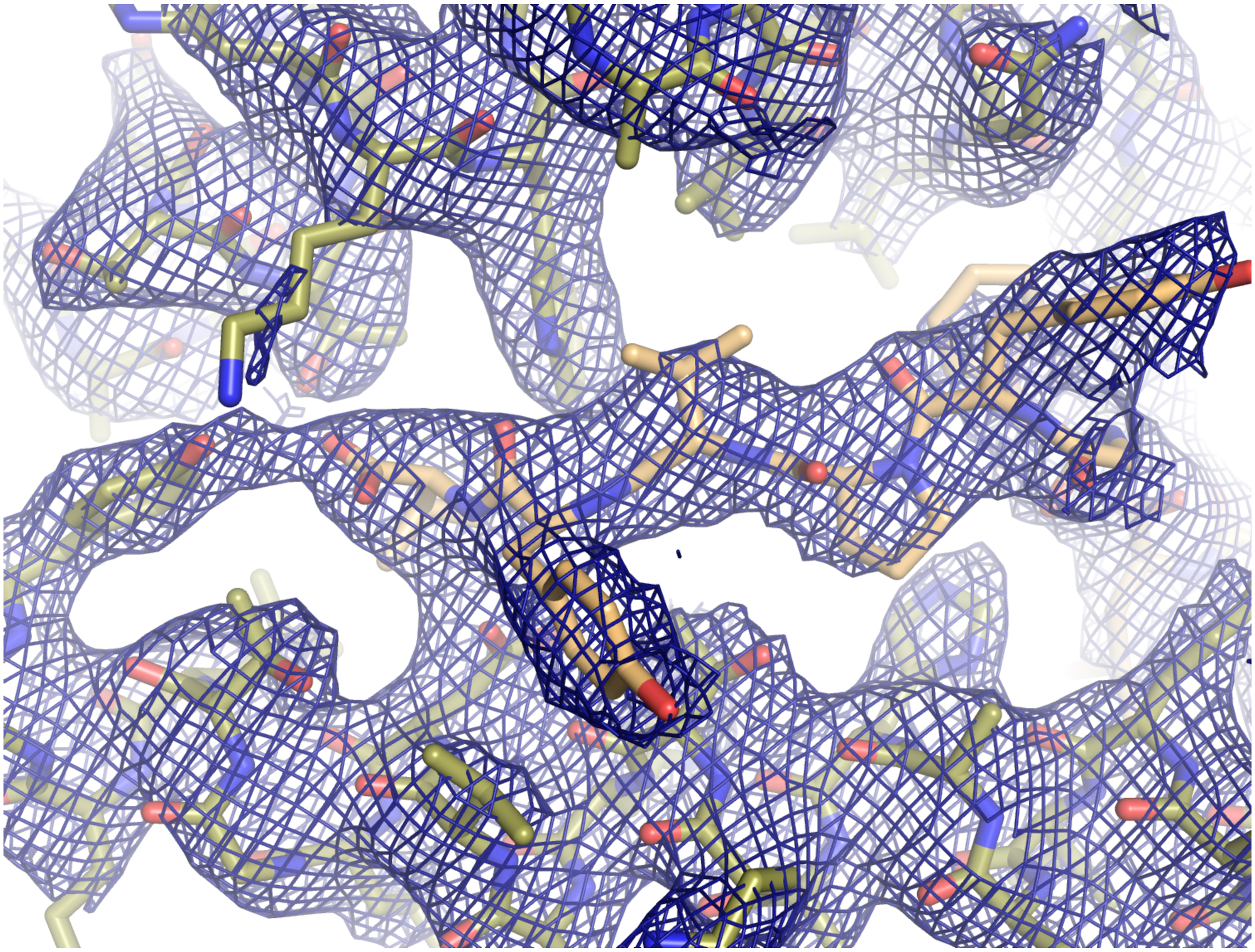

**Figure S6. Detail of electron density around TAX9 peptide.** 2mFo-DFc map (blue) contoured at 1 σ around TAX9 peptide (light orange) and CSM8 A*02:01 helices α2 (top, olive) and α1 (bottom).

**Supplementary Table S1. Crystallographic data collection and refinement statistics for CSM8 H-2D$^{b}$/ gp33**

| | |
|---|---|
| PDB | 9HY4 |
| Space group | C 1 2 1 |
| Cell dimensions | |
| *a* (Å) | 65.95 |
| *b* (Å) | 93.24 |
| *c* (Å) | 75.86 |
| b (°) | 94.08 |
| Resolution (Å) | 53.75-2.00 (2.05-2.00) |
| $N_{unique}$ | 29279 (2209) |
| Multiplicity | 2.2 (2.3) |
| Completeness (%) | 95.2 (96.9) |
| I/s(I) | 9.4 (4.1) |
| $R_{merge}$ | 0.057 (0.103) |
| Complexes in au | 2 |
| Refinement | |
| $R_{cryst}$ (%) | 23 |
| $R_{free}$ (%) | 28 |
| Rmsd from ideal geometry | |
| Bond length (Å) | 0.009 |
| Bond angles (°) | 1.049 |
| Ramachandran plot | |
| Residues in preferred regions (%) | 97.42 |
| Outliers (%) | 0 |

Rmerge=Σ|Ii−Im| / Σ Ii, where Ii is the intensity of the measured reflection, and Im is the mean intensity for all observations of that reflection. Numbers within parentheses are for the outer resolution shell of data.

**Supplemental Table S2. Crystallographic data collection and refinement statistics for A6c134/CSM8 A*02/TAX9.**

| | |
|---|---|
| PDB accession code | 9NDS |
| Wavelength (Å) | 0.9202 |
| Resolution range (Å) | 46.62 - 3.4 (3.52 - 3.4) |
| Space group | P 31 |
| Unit cell (Å, °) | 106.906 106.906 190.592 90 90 120 |
| Total reflections | 117680 (12091) |
| Unique reflections | 33320 (3374) |
| Multiplicity | 3.5 (3.6) |
| Completeness (%) | 99.25 (99.29) |
| Mean I/sigma(I) | 4.33 (0.64) |
| Wilson B-factor (Å$^2$) | 107.63 |
| R-merge | 0.2782 (2.599) |
| R-meas | 0.3274 (3.046) |
| R-pim | 0.1707 (1.576) |
| CC1/2 | 0.976 (0.149) |
| Reflections used in refinement | 33276 (3353) |
| Reflections used for R-free | 1377 (142) |
| R-work | 0.2173 (0.3478) |
| R-free | 0.2683 (0.3903) |
| Number of non-hydrogen atoms | 10960 |
| macromolecules | 10866 |
| ligands | 94 |
| solvent | 0 |
| Protein residues | 1370 |
| RMS(bonds) (Å) | 0.002 |
| RMS(angles) (°) | 0.47 |
| Ramachandran favored (%) | 96.81 |
| Ramachandran outliers (%) | 0.00 |
| Rotamer outliers (%) | 3.26 |
| Clashscore | 4.47 |
| Average B-factor (Å$^2$) | 125.67 |
| macromolecules | 124.86 |
| ligands | 219.59 |

Statistics for the highest-resolution shell are shown in parentheses.

**Supplemental table S3. Stabilizing domain amino acid sequences.**

| Name | Sequence |
|---|---|
| hit6 stabilizer | DREDVERLLRAVEWAIKAGDPYSARVLVELAREDAEKIGDERLRREVEELLRELEELGGSGGSGGSGGSGGSGG |
| CSM8 stabilizer | DREDVERLLRSVEWAIKAGDPYSARILVELAREDAEKIGDERLRREVEELLRELEELGGSGGSGGSGGSGGSGG |
| SMART stabilizer | DREDVERLLRSVEWAIKAGDPYSARILVELAREDAEKIGDERLRREVEELLRELEELEEELARLPKLPPD |

**Supplemental table S4. MHC peptide binding groove amino acid sequences (α1+α2 domains).** All sequences shown below contain the CSM8 mutations unless otherwise noted.

| Name | Sequence |
|---|---|
| H-2D$^{b}$ (WT) | GPHSMRYFETAVSRPGLEEPRYISVGYVDNKEFVRFDSDAENPRYEPRAPWMEQEGPEYWERETQKAKGQEQWFRVSLRNLLGYYNQSAGGSHTLQQMSGCDLGSDWRLLRGYLQFAYEGRDYIALNEDLKTWTAADMAAQITRRKWEQSGAAEHYKAYLEGECVEWLHRYLKNGNATL |
| H-2D$^{b}$ | GPHSMKYIETAISRPGLEEPRYISVGYVDNKEFVRFDSDAENPRYEPRAPWMEQEGPEYWERETQKAKGQEQWFRVSLRNLLGYYNQSAGGSHTLQQMSGCDLDENWRLVRGYLQFAYEGRDYIALNEDLKTWTAADMAAQITRRKWEQSGAAEHYKAYLEGECVEWLHRYLKNGNATL |
| H-2D$^{b}$ (WT + Y84A) | |
| A*02:01 (WT) | GSHSMRYFFTSVSRPGRGEPRFIAVGYVDDTQFVRFDSDAASQRMEPRAPWIEQEGPEYWDGETRKVKAHSQTHRVDLGTLRGYYNQSEAGSHTVQRMYGCDVGSDWRFLRGYHQYAYDGKDYIALKEDLRSWTAADMAAQTTKHKWEAAHVAEQLRAYLEGTCVEWLRRYLENGKETLQ |
| A*02:01 | GSHSMKYIFTSISRPGRGEPRFIAVGYVDDTQFVRFDSDAASQRMEPRAPWIEQEGPEYWDGETRKVKAHSQTHRVDLGTLRGYYNQSEAGSHTVQRMYGCDVDENWRFVRGYHQYAYDGKDYIALKEDLRSWTAADMAAQTTKHKWEAAHVAEQLRAYLEGTCVEWLRRYLENGKETLQ |
| A*02:01 (WT + W167A) | GSHSMRYFFTSVSRPGRGEPRFIAVGYVDDTQFVRFDSDAASQRMEPRAPWIEQEGPEYWDGETRKVKAHSQTHRVDLGTLRGYYNQSEAGSHTVQRMYGCDVGSDWRFLRGYHQYAYDGKDYIALKEDLRSWTAADMAAQTTKHKWEAAHVAEQLRAYLEGTCVEALRRYLENGKETLQR |
| A*01:01 | GSHSMKYIFTSISRPGRGEPRFIAVGYVDDTQFVRFDSDAASQKMEPRAPWIEQEGPEYWDQETRNMKAHSQTDRANLGTLRGYYNQSEDGSHTIQIMYGCDVDENGRFVRGYRQDAYDGKDYIALNEDLRSWTAADMAAQITKRKWEAVHAAEQRRVYLEGRCVDGLRRYLENGKETLQ |

| | |
|---|---|
| A*03:01 | GSHSMKYIFTSISRPGRGEPRFIAVGYVDDTQFVRFDSDAASQRMEPRAPWIEQEGPEYWDQETRNVKAQSQTDRVDLGTLRGYYNQSEAGSHTIQIMYGCDVDENGRFVRGYRQDAYDGKDYIALNEDLRSWTAADMAAQITKRKWEAAHEAEQLRAYLDGTCVEWLRRYLENGKETLQ |
| A*24:02 | GSHSMKYISTSISRPGRGEPRFIAVGYVDDTQFVRFDSDAASQRMEPRAPWIEQEGPEYWDEETGKVKAHSQTDRENLRIALRYYNQSEAGSHTLQMMFGCDVDENGRFVRGYHQYAYDGKDYIALKEDLRSWTAADMAAQITKRKWEAAHVAEQQRAYLEGTCVDGLRRYLENGKETLQ |
| A*29:02 | GSHSMKYITTSISRPGRGEPRFIAVGYVDDTQFVRFDSDAASQRMEPRAPWIEQEGPEYWDLQTRNVKAQSQTDRANLGTLRGYYNQSEAGSHTIQMMYGCDVDENGRFVRGYRQDAYDGKDYIALNEDLRSWTAADMAAQITQRKWEAARVAEQLRAYLEGTCVEWLRRYLENGKETLQ |
| A*30:01 | GSHSMKYISTSISRPGSGEPRFIAVGYVDDTQFVRFDSDAASQRMEPRAPWIEQERPEYWDQETRNVKAQSQTDRVDLGTLRGYYNQSEAGSHTIQIMYGCDVDENGRFVRGYEQHAYDGKDYIALNEDLRSWTAADMAAQITQRKWEAARWAEQLRAYLEGTCVEWLRRYLENGKETLQ |
| B*07:02 | GSHSMKYIYTSISRPGRGEPRFISVGYVDDTQFVRFDSDAASPREEPRAPWIEQEGPEYWDRNTQIYKAQAQTDRESLRNLRGYYNQSEAGSHTLQSMYGCDVDENGRLVRGHDQYAYDGKDYIALNEDLRSWTAADTAAQITQRKWEAAREAEQRRAYLEGECVEWLRRYLENGKDKLE |
| B*08:01 | GSHSMKYIDTAISRPGRGEPRFISVGYVDDTQFVRFDSDAASPREEPRAPWIEQEGPEYWDRNTQIFKTNTQTDRESLRNLRGYYNQSEAGSHTLQSMYGCDVDENGRLVRGHNQYAYDGKDYIALNEDLRSWTAADTAAQITQRKWEAARVAEQDRAYLEGTCVEWLRRYLENGKDTLE |
| B*15:01 | GSHSMKYIYTAISRPGRGEPRFIAVGYVDDTQFVRFDSDAASPRMAPRAPWIEQEGPEYWDRETQISKTNTQTYRESLRNLRGYYNQSEAGSHTLQRMYGCDVDENGRLVRGHDQSAYDGKDYIALNEDLSSWTAADTAAQITQRKWEAAREAEQWRAYLEGLCVEWLRRYLENGKETLQ |
| B*37:01 | GSHSMKYIHTSISRPGRGEPRFISVGYVDDTQFVRFDSDAASPRTEPRAPWIEQEGPEYWDRETQISKTNTQTYREDLRTLLRYYNQSEAGSHTIQRMSGCDVDENGRLVRGYNQFAYDGKDYIALNEDLSSWTAADTAAQITQRKWEAARVAEQDRAYLEGTCVEWLRRYLENGKETLQ |
| B*38:01 | GSHSMKYIYTSISRPGRGEPRFISVGYVDDTQFVRFDSDAASPREEPRAPWIEQEGPEYWDRNTQISKTNTQTYRENLRIALRYYNQSEAGSHTLQRMYGCDVDENGRLVRGHNQFAYDGKDYIALNEDLSSWTAADTAAQITQRKWEAARVAEQLRTYLEGTCVEWLRRYLENGKETLQ |
| B*58:01 | GSHSMKYIYTAISRPGRGEPRFIAVGYVDDTQFVRFDSDAASPRTEPRAPWIEQEGPEYWDGETRNMKASAQTYRENLRIALRYYNQSEAGSHIIQRMYGCDLDENGRLVRGHDQSAYDGKDYIALNEDLSSWTAADTAAQITQRKWEAARVAEQLRAYLEGLCVEWLRRYLENGKETLQ |
| E*01:03 | GSHSLKYIHTSISRPGRGEPRFISVGYVDDTQFVRFDNDAASPRMVPRAPWM |

| | |
|---|---|
| | EQEGSEYWDRETRSARDTAQIFRVNLRTLRGYYNQSEAGSHTLQWMHGCEL DENGRFVRGYEQFAYDGKDYLTLNEDLRSWTAVDTAAQISEQKSNDASEAEH QRAYLEDTCVEWLHKYLEKGKETLL |
| F*01:01 | GSHSLKYISTAISRPGRGEPRYIAVEYVDDTQFLRFDSDAAIPRMEPREPWVEQ EGPQYWEWTTGYAKANAQTDRVALRNLLRRYNQSEAGSHTLQGMNGCDMD ENGRLVRGYHQHAYDGKDYISLNEDLRSWTAADTVAQITQRFYEAEEYAEEFR TYLEGECLELLRRYLENGKETLQ |
| G*01:01 | GSHSMKYISAAISRPGRGEPRFIAMGYVDDTQFVRFDSDSASPRMEPRAPWV EQEGPEYWEEETRNTKAHAQTDRMNLQTLRGYYNQSEASSHTLQWMIGCDL DENGRLVRGYERYAYDGKDYLALNEDLRSWTAADTAAQISKRKSEAANVAEQR RAYLEGTCVEWLHRYLENGKEMLQ |

**Supplemental table S5. Peptide amino acid sequences.**

| **Name** | **Sequence** |
|---|---|
| gp33 | KAVYNFATM |
| gp33 V3P | KAPYNFATM |
| gp33 M9C | KAVYNFATC |
| gp33 PF | KAPFNFATM |
| gp33 Y4F | KAVFNFATM |
| TAX9 | LLFGYPVYV |
| NYESO-1 | SLLMWITQC |
| NYESO-1 9V | SLLMWITQV |
| Mart1 | AAGIGILTV |

**Supplemental table S6. Other amino acid sequences.**

| **Name** | **Sequence** |
|---|---|
| Aga2 (with signal peptide) | MQLLRCFSIFSVIASVLAQELTTICEQIPSPTLESTPYSLSTTTILANG KAMQGVFEYYKSVTFVSNCGSHPSTTSKGSPINTQYVF |
| Aga2 linker (with HA tag) | KDNSSTIEGRTRGSGSGSYPYDVPDYAGS |
| Peptide linker (yeast display) | GGGGSGGGGSGGGGS |
| SUMO | MDSEVNQEAKPEVKPEVKPETHINLKVSDGSSEIFFKIKKTTPLRRL MEAFAKRQGKEMDSLRFLYDGIRIQADQAPEDLDMEDNDIIEAHRE |

| | |
|---|---|
| | QIGG |
| Peptide linker (oligomer) | GGGASGGGGSGGGGS |
| $Tet_{T=4}$-4 (oligomer) | PELFLQDLRSLVEAARILARLARQRGDEHALERAARWAEQAARQAERLARQARKEGNLELALKALQILVNAAYVLAEIARDRGNEELLEYAARLAEEAARQAIEIWANAMIEGNQQLRTKAAHIILRAAEVLLEIARDRGNQELLEKAASLVDAVAALQQAAAAILEGDVEKAVRAAQEAVKAAKEAGDNDMLNAVASAAARIALEAMQQGNLEVALKALRVAEEALKQAGGSGGSHHHHHH |
| Oligomer linker | GGSGGSGGSGGSGGSGGSGGSGGSGGSGS |
| Myc tag (with linker) | GGSGGSGGSSEQKLISEEDL |
| Ulp1 | MGGLVPELNEKDDDQVQKALASRENTQLMNRDNIEITVRDFKTLAPRRWLNDTIIEFFMKYIEKSTPNTVAFNSFFYTNLSERGYQGVRRWMKRKKTQIDKLDKIFTPINLNQSHWALGIIDLKKKTIGYVDSLSNGPNAMSFAILTDLQKYVMEESKHTIGEDFDLIHLDCPQQPNGYDCGIYVCMNTLYGSADAPLDFDYKDAIRMRRFIAHLILTDALKG |

*Sequence construction*
SMART MHC variants can be constructed by directly fusing sequences as follows:

**Monomer:** [stabilizer; table S3] + [MHC; table S4]
We recommend using the CSM8 mutant version of the MHC when pairing with the CSM8 or SMART stabilizer.

**Yeast Display:** [Aga2; table S6] + [Aga2 linker; table S6] + [stabilizer; table S3] + [MHC; table S4] (+ [peptide linker (yeast display); table S6] + [peptide; table S5])
Peptide display is optional, but if the peptide is included, the W167A mutation should be included in the MHC.

**Oligomer:** [SUMO; table S6] + [peptide; table S5] + [peptide linker (oligomer); table S6] + [stabilizer; table S3] + [MHC (with Y84A); table S4] + [oligomer linker; table S6] + [$Tet_{T=4}$-4; table S6]
We recommend using the SMART stabilizer and CSM8 mutations in oligomeric designs. These should be co-expressed with Ulp1 (table S6) to cleave the SUMO tag.
